# Oldest attested languages in the Near East reveal deep transformations in the distribution of linguistic features

**DOI:** 10.1101/2024.06.25.600575

**Authors:** Nour Efrat-Kowalsky, Peter Ranacher, Nico Neureiter, Paul Widmer, Balthasar Bickel

## Abstract

It is an unresolved question to what extent the current frequency distributions of linguistic features inform us about what is representative of the language faculty and does not instead result from historical contingencies. We probe this question by leveraging unique data from the oldest attested languages, those preserved through writing from up to 5,000 years ago in the Ancient Near East. We examine 70 grammatical features for which there is sufficient evidence in the available records. After controlling for relatedness we find robust deviations of two of the oldest languages, Hurrian and Sumerian from both the ancient languages and the modern distribution. The spatial and temporal placement of these two languages reveal a divergent distribution of features in the region in prehistory, suggesting massive transformations of the linguistic distributions in the past few millennia. This challenges inferences about general characteristics of language based on modern distributions.

## 1 Introduction

Characterizations of the language faculty are often based on frequencies of specific linguistic features (for example word order), assuming that the asymmetries in these frequencies reveal universal principles of language or of general cognitive and communicative processes [1–3]. However, what we know of those frequencies is based on the current distribution of the world’s languages. Yet, current distributions are subject to a major caveat. It is quite possible that a currently dominant feature was dispreferred some time in the distant past [4–7], reflecting local population history rather than a universal preference [8–11].

Indeed, population history has often changed the distribution of languages, leading to regular processes of language shift [12]. This has erased much of earlier linguistic diversity and thereby re-shaped the frequency distribution of linguistic features. In the Eurasian steppe, for example, the spreads of Iranian, Turkic and Mongolic speakers have erased nearly all of the indigenous languages of Europe (Basque being the only one remaining), the languages of the Trypillia-Cucuteni culture, Avar (known by name only), languages of the North Caucasian plain, the languages of the Bactria-Margiana Archaeological Complex, the Iranian Scythian, the Germanic Gothic, Hunnish (probably Bulgar Turkic), Kitan and other Para-Mongolic languages [13]. In Africa, the spread of Berber and Arabic in the north, and Bantu in Sub-Saharan Africa have caused many indigenous languages to go extinct and reduced the diversity within their families [14]. Language spreads are also documented in antiquity. In the Near East, Sumerian was replaced by Akkadian, which was later replaced by Aramaic, which was replaced by Arabic. In Europe, the Italic spread replaced local languages such as Etruscan and Continental Celtic. Language spreads may lead to the extinction of the local languages through language shift caused by the assimilation of the local population or through the dying out of the local population due to genocide and/or new diseases brought in with the spread [15].

We are still missing the full picture on the structural diversity that was wiped out. To assess the extent to which earlier distributions might have been different from present-day distributions of linguistic featires, we focus on the Ancient Near East (ANEA), a region in which we have access to the oldest documentation of linguistic structural features.

The ANEA was a linguistically diverse region, spanning over millennia, with languages from unrelated genealogies. Though comparative historical linguistics can give some insight into unattested linguistic prehistory, it is limited to reconstructions within a given family of historically related languages (such as Indo-European or Semitic) and cannot access prehistoric distributions across families and entire regions [16–19]. We asses the extent to which the frequency distributions of linguistic features in the ANEA differed from the modern distribution with two sets of analyses, one comparing the ANEA as a whole with the current global distribution outside the ANEA in a regression framework, one exploring possible sub-clusters within the ANEA with what is expected from global or family-specific patterns in a clustering framework. We hypothesized that a statistically robust difference at any of these levels is indicative of change in frequency distributions in the past 5,000 years - the time of the earliest known documents in the ANEA. Alternatively, if there is no appreciable difference, this would suggest that frequency distributions are independent of time, possibly reflecting stationary (equilibrium) probabilities [4, 20]. To test these hypotheses we created a database of 70 grammatical features for which the available record was sufficiently reliable (Supplementary Information, Section S1).

## 2 Methods

### 2.1 Data

During the time from the mid-fourth millennium BCE until the mid-first millennium CE we know of cultures living in the Near East speaking at least 17 languages from at least six different phylogenetic clades including: Sumerian, Elamite, Hurrian-Urartian (all isolates with no established genetic relationship to other known languages^1^), Akkadian, Eblaite, Amorite, Aramaic, Ugaritic, Phoenician, Hebrew, Ancient North Arabian, Ancient South Arabian, Arabic (Semitic), Egyptian, Coptic (Egyptian), Hittite, and Luwian (Indo-European) [21] (Figure 1).

**Figure 1:**
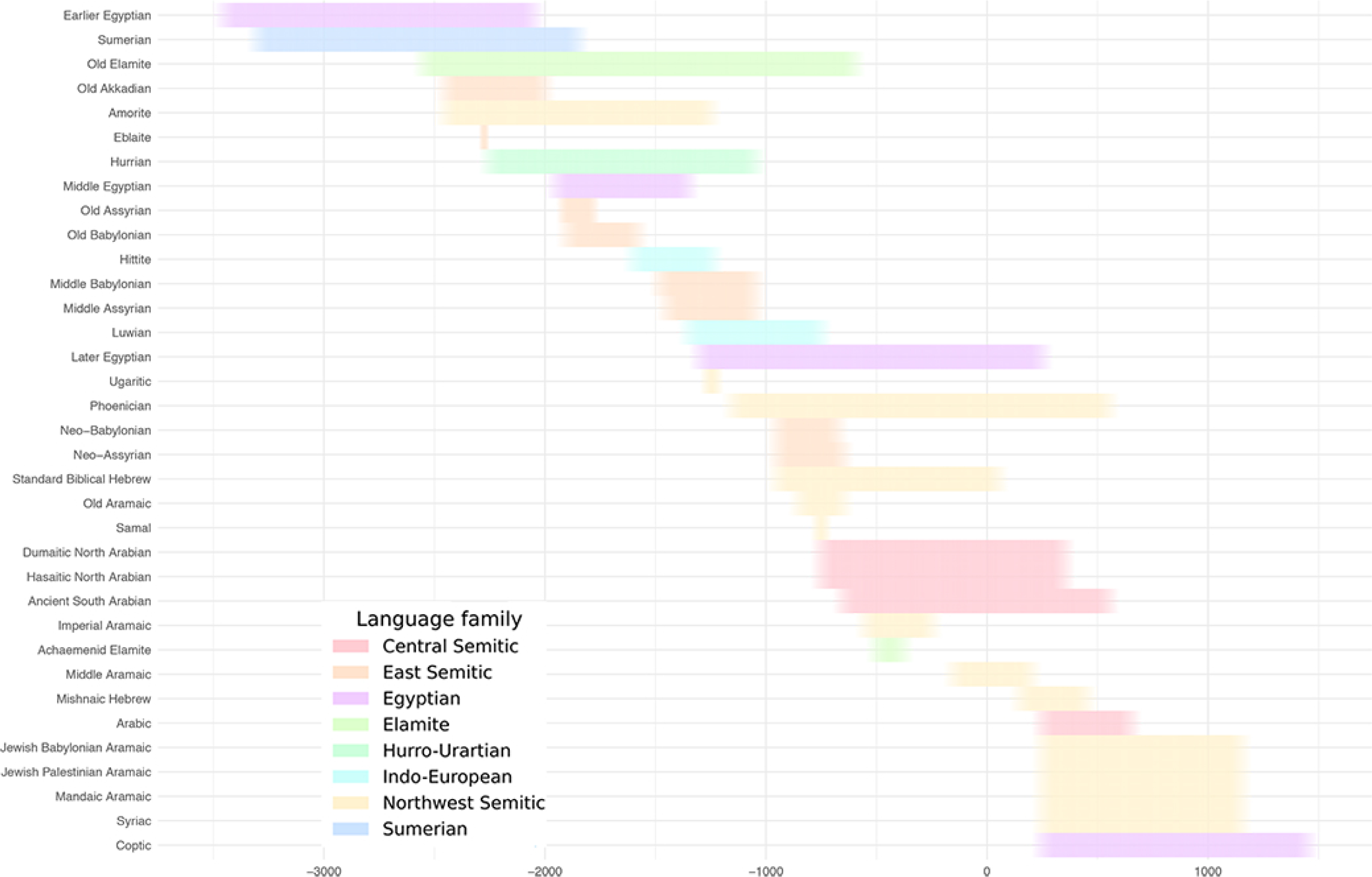
The language varieties in the Ancient Near East sample by date. Dates refer to time of attestation.

The data for the languages of the Ancient Near East sample were collected from grammars and contains 35 distinct language varieties, which either represent a language, a distinct local dialect of a language (local variety), or a distinct diachronic period of a language (diachronic variety). The full list of language varieties and the data sources for each is given in Table S2 of the Supplementary Information. The attestation of some language varieties is more complete than others. If a feature value could not be determined due to too meagre attestation, this data point was left blank (see below for how we deal with missing values in each analysis). Where two grammars of the same language variety gave a conflicting analysis for a feature, we made choices based on the examples provided in the grammars, though such cases were exceedingly rare.

Feature selection followed the principle that no feature is exempt in principle from the effects of language change [22–26]. Data was collected on all primary morphological and syntactic features from [27] and [28] that are present in the Ancient Near East languages. Altogether, data was collected for 51 original features and recoded into 70 in order to resolve logical dependencies between features. This resulted in a total of 192 feature states. A list of all the features used with their abbreviated names and the corresponding names in WALS and AUTOTYP is given in Table S1 of the Supporting Information. Duplicate features that appear in both databases were combined. We selected features from published typological databases, rather than focusing on features that are only relevant to the ANEA such that we can compare the ANEA data to data from a representative sample of language from around the world. Phonetic, phonological and semantic features were not included in the dataset. The Ancient Near Eastern data is extracted from written attestations of dead languages with no living speakers or recordings available. Any claim about the phonology, sound inventories or specific semantic distinctions is inherently insecure and subject to reconstructions. Similarly, semantic interpretations are difficult to verify in the absence of native speakers. Features from morphology and syntax are more overtly visible in the texts and are normally straightforward to analyze.

To find out if the ANEA languages are distinct from current linguistic distributions and its succeeding language families, we contrasted the ANEA data with data from a sample of 100 languages from around the world. This sample is balanced in terms of areal and family distributions [27]. The data was downloaded from the and databases and accessed through the lingtypology R package [29]. The 100 languages in our sample are the languages with the least amount of missing data with regard to the features we were targeting.

Although within-language variation (for example variable word order in a language) is an important attribute of language, it is not captured in the available data with sufficient systematicity. In all cases of known variation we entered the default, basic or most frequent variant into our database.

### 2.2 Regression

In order to test the hypothesis that the Ancient Near East has a distinctive frequency distribution from the rest of the world, i.e. current languages, we compared the ANEA language sample to the universal language sample. We applied logistic and categorical (multinomial) Bayesian regression for the binary and multi-state features, respectively. The grammatical feature is the dependent variable, the area is the predictor variable, and controlling for language family as random effects. Area here refers to a language being in or outside the Ancient Near East. The models estimate the posterior probability of an effect of area on a feature.

We chose *N* (0, 1) as the prior for the area predictor, which is relatively uniform on the response scale [30]. For the group-level (“random”) effects, we used a Student-*t* (3,0,2.5) prior. In a sensitivity analysis we found that more uninformative priors do not substantially alter our results. For fitting the models, we used the R package brms as an interface to Stan [31, 32]. Missing values were excluded from the models.

### 2.3 Clustering: sBayes

For clustering we use sBayes [33], a Bayesian mixture model that infers clusters of languages through patterns that can neither be attributed to family membership (i.e. shared descent) nor to global preferences. The advantage of this model over the regression is that it allows discovery of sub-clusters within the ANEA. In other words, while the regression can only test whether the ANEA as a whole differs from the global distributions, sBayes allows discovery of specific *clusters* within the ANEA that differ from the global distributions.

Specifically, sBayes achieves this through a generative model that explains the data by a mixture of three categorical distributions: a universal likelihood (*α*), a family likelihood (*β*) and a cluster likelihood (*γ*). The universal likelihood affects all languages equally. In contrast, the family likelihood differs between language families, and the cluster likelihood differs between the clusters that the method ultimately aims to find. Each of these three components has corresponding weights – *w_universal_*, *w_family_*and *w_cluster_*– which can vary between the different features. The likelihood of observing state *s* for feature *f* in language *l* is defined by:

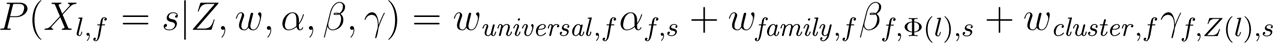

Here, *Z* is the assignment of languages to clusters, and Φ is the assignment of languages to language families. sBayes takes the family assignment Φ as a given. Hence, we omit it on the left-hand side of the equation. The overall likelihood of a data set *D* is the product over the likelihood of all features and languages:

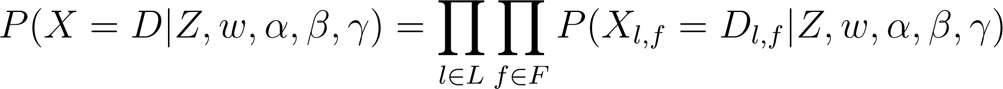

If a feature value is missing for a language, the corresponding term in this product will be 1 and, hence, the missing value will not influence the likelihood.

The sBayes model relies on priors for each parameter (*Z, w, α, β, γ*). For *Z*, *w* and *γ* we use a uniform prior [34], since we do not want to impose any prior expectation about cluster membership (*Z*), the importance of different mixture components (*w*) or the distributions in the clusters (*γ*). For *α* (universal) and *β* (family), we impose empirical priors based on observations outside our sample.

For *α*, we use an empirical prior on the basis of our global 100-languages sample, captured by a Dirichlet distribution:

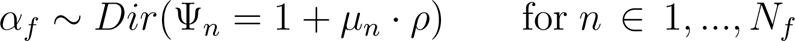

Here, *N_f_* is the number of states in feature *f*. *µ_n_* is the frequency of state *n* in the global sample and defines the mean of the prior distribution. *ρ* gives the precision or inverse variance of the Dirichlet distribution. Ψ*_n_* can be considered pseudocounts for state *n*, i.e. counts corrected by uncertainty and ensuring that each state remains a possibility (hence +1). A large precision yields large pseudocounts and a narrow prior. We vary the precision *ρ* for the universal prior distribution (from *ρ* = 10 to *ρ* = 90) in sensitivity analyses (Section S3.1.1., Supplementary Information).

For *β*, we informed the family distribution prior with data for the Semitic languages. All three branches of the Semitic language family in our sample were informed by the same family distribution which included data for a sample of the whole of the Semitic language family outside our sample. The data informing the family distribution for the Semitic languages was accessed in the same way as the data for the universal distribution. It was supplemented by data collected from grammars for languages and branches that were not represented in the database to ensure a more balanced sample. We set the precision *ρ* for the family prior distribution low (*ρ* = 10) for the same reasons as were given for the universal prior on *α*.

### 2.4 Data availability

The data and script to run the analyses in this paper are available on https://github.com/noorefrat/Oldest-attested-languages-in-the-Near-East.

## 3 Results

In the regression analysis the masses of the posterior probability of an area effect concentrate around 0 for most features, suggesting no clear difference in the (log) odds for features inside vs. outside the Ancient Near East. Only 7 out of 192 feature-states have 90% of the highest probability density estimates above or below 0. This could either reflect low power of the analysis or large similarity between the ANEA languages as a whole and the universal language sample, baring in mind that in such a large sample it is statistically expected that a handfull would show an effect (Figure 2).

**Figure 2:**
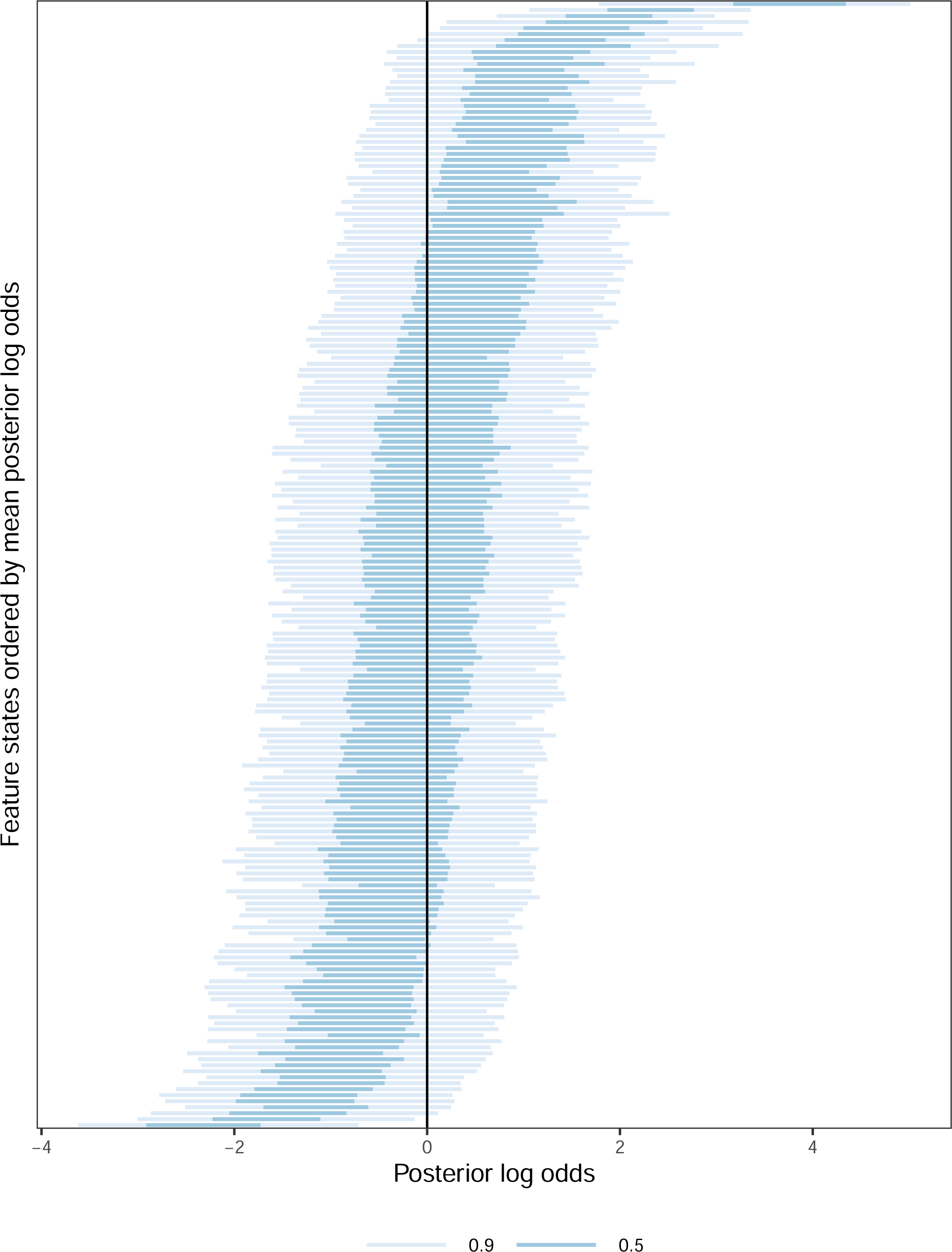
Posterior distributions of the predictor that marks ANEA languages for all grammatical feature states. For all but 7 feature states, the posterior probabilities are concentrated on log odds at or near zero which is marked with a black vertical line, suggesting no difference in the structural profiles between languages inside and languages outside the Ancient Near East (See Fig. S1 and S2 for the same plot with explicit feature-state labels)

The sBayes model resulted in one cluster containing two languages: Hurrian and Sumerian (Figure 3). The other languages of the Ancient Near East were not assigned to any cluster, meaning that their similarities were better explained by the universal or family-specific distributions. The quantitative evidence is consistent with qualitative inspection of individual features where the dominant feature states in the Ancient Near East tend to replicate the universal norm with few exceptions. The features that had the highest weight (see S3.2.4) in clustering Hurrian and Sumerian against all other languages are a) marking of plural nouns, where the most common marking worldwide, plural suffix, is also most common in the Ancient Near East languages, with the exception of Sumerian which marks the plural with a clitic; b) marking of negation, where the most common marking worldwide, negation particle, is also most common in the Ancient Near East languages, with the exception of Sumerian and Hurrian which mark negation with a variation between negative word and affix and a negative affix respectively; and c) number of inflectional categories: low number of inflectional categories is more common worldwide and in the Ancient Near East, but both Sumerian and Hurrian exhibit a higher number of inflectional categories (Figure 4).

**Figure 3:**
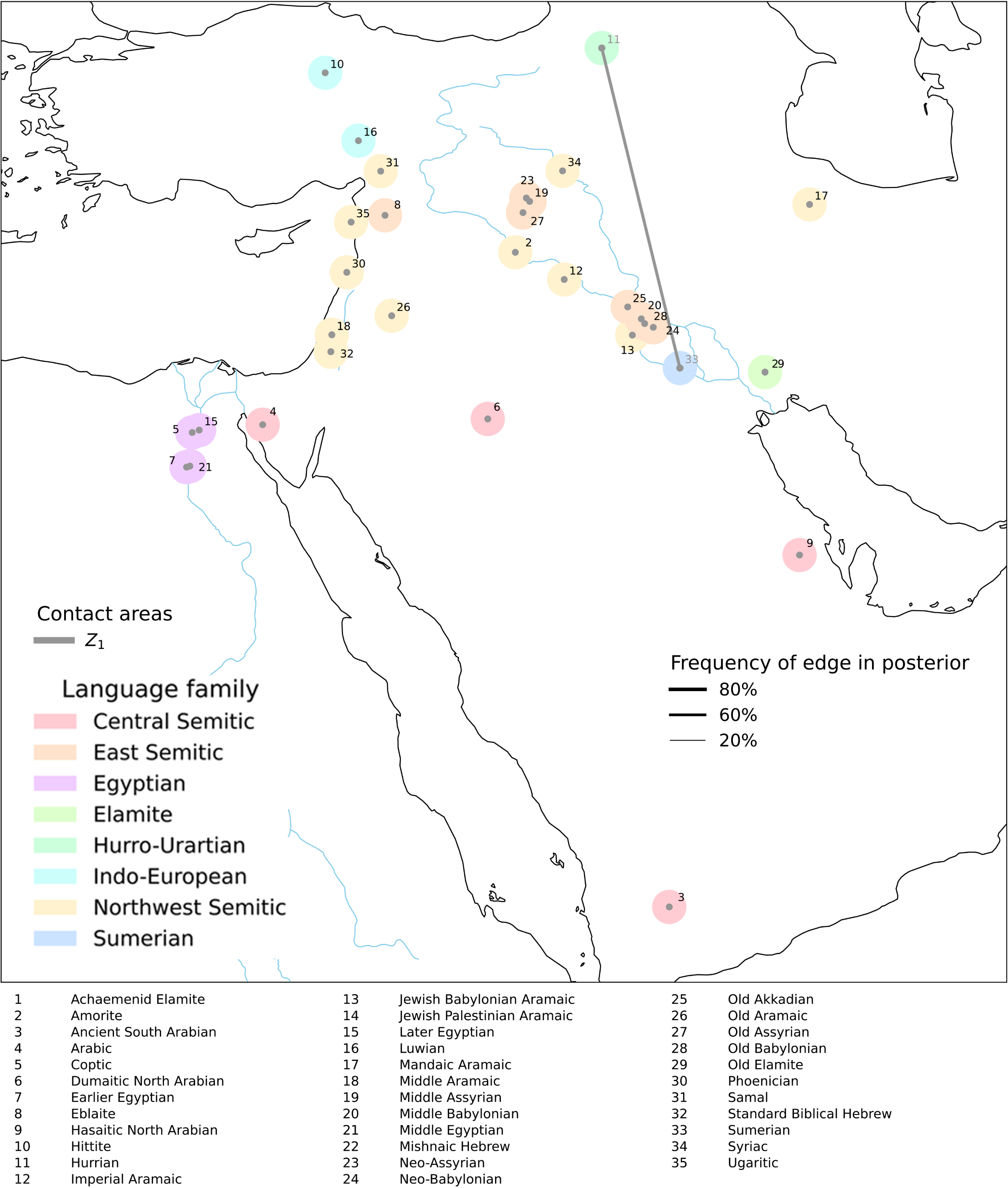
The Hurrian-Sumerian cluster. A mixture model reveals a high posterior probability that Hurrian and Sumerian, but no other languages, share feature states unexplained by phylogenetic and universal trends. All the languages in the dataset are colored according to their phylogenetic affiliation. Clusters are marked by a line connecting the languages (here, Hurrian and Sumerian). The evidence for the cluster is very strong, with the two languages appearing together in the posterior distribution in at least 80% of samples (Section S3.2.1, Supplementary Information)

**Figure 4:**
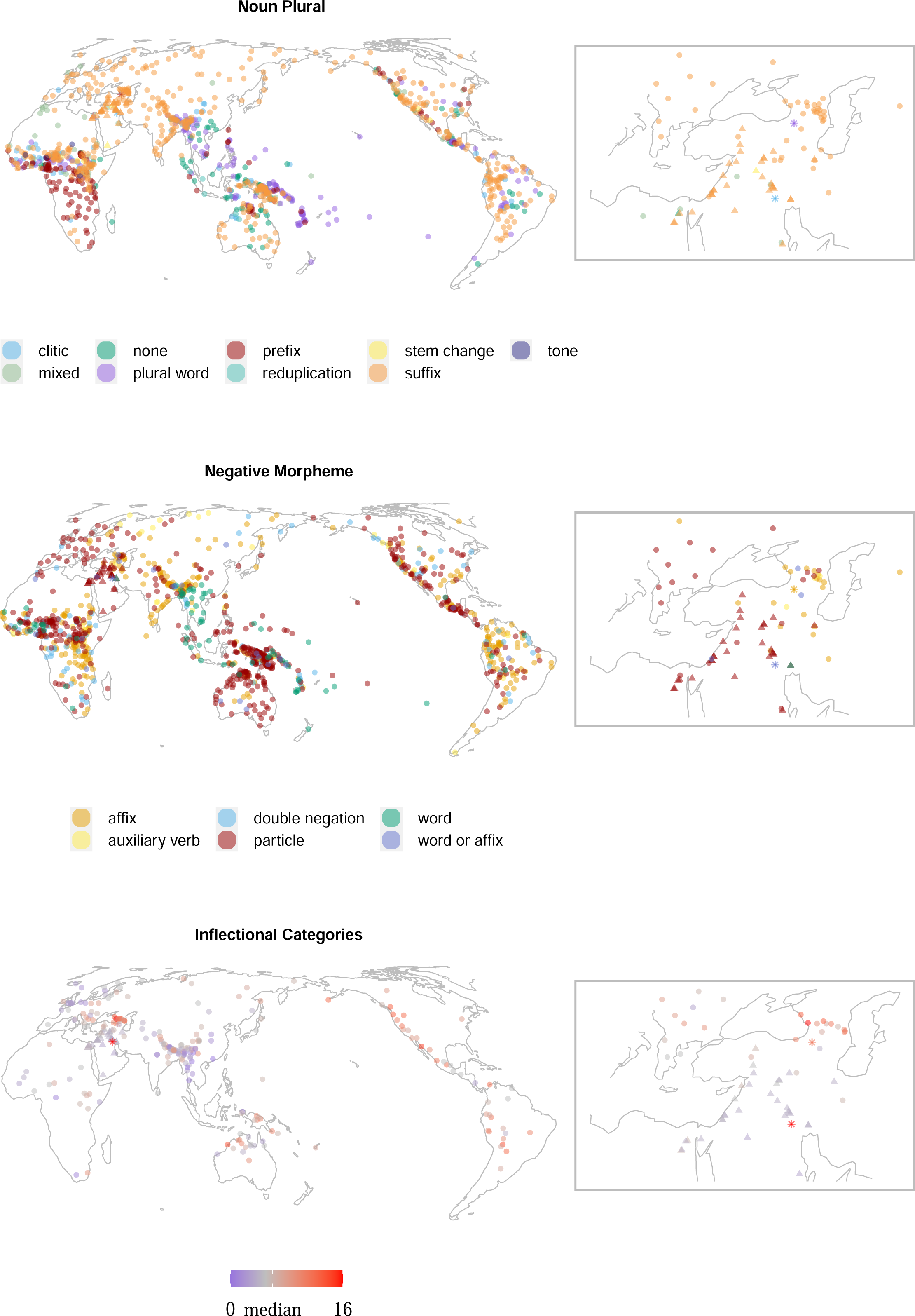
Representative distributions of the features Noun Plural Marking, Negative Morpheme, and Number of Inflectional Categories. Ancient Near East languages are marked by a triangle. Hurrian and Sumerian are marked by a star. Inset on the right on the Near East.

We performed a series of sensitivity analyses to verify the robustness of the signal (Section S3.1, Supplementary Information). The majority of the language varieties in our sample are Semitic. We therefore manipulated the amount of information on phylogenetic relatedness. In the main analysis, we captured relatedness at the level of major branches. Correspondingly, the Semitic languages are considered as three different language families at the major-branch level, with no further relation between them: East Semitic, Central Semitic, and Northwest Semitic [35]. To maximize the detectability of clusters, we also fitted a model without information on family relations, i.e. without a mixture component for the (Semitic) families. The results of this analysis again only suggested the Hurrian and Sumerian cluster. In this version of the model, where inheritance in a family was taken out as an explanatory factor for language similarities, all other similarities between the languages were best explained by universal preferences (Section S3.1.2, Supplementary Information). This result is especially striking because of the high similarity between the Semitic languages. Even though the Semitic languages in our sample span nearly 3,000 years, there is remarkably little variation over time in the values of the features (with a mean consistency of .69; Figure 5). This shows that the Semitic languages are quite similar to one another with regard to the morphosyntactic features tested here, regardless of when and where they were spoken. In the absence of the family component in the model, the low entropy exhibited by the Semitic languages would be expected to override the similarity of the Semitic languages to the universal tendencies and be assigned to clusters, yet this still isn’t the case.

**Figure 5:**
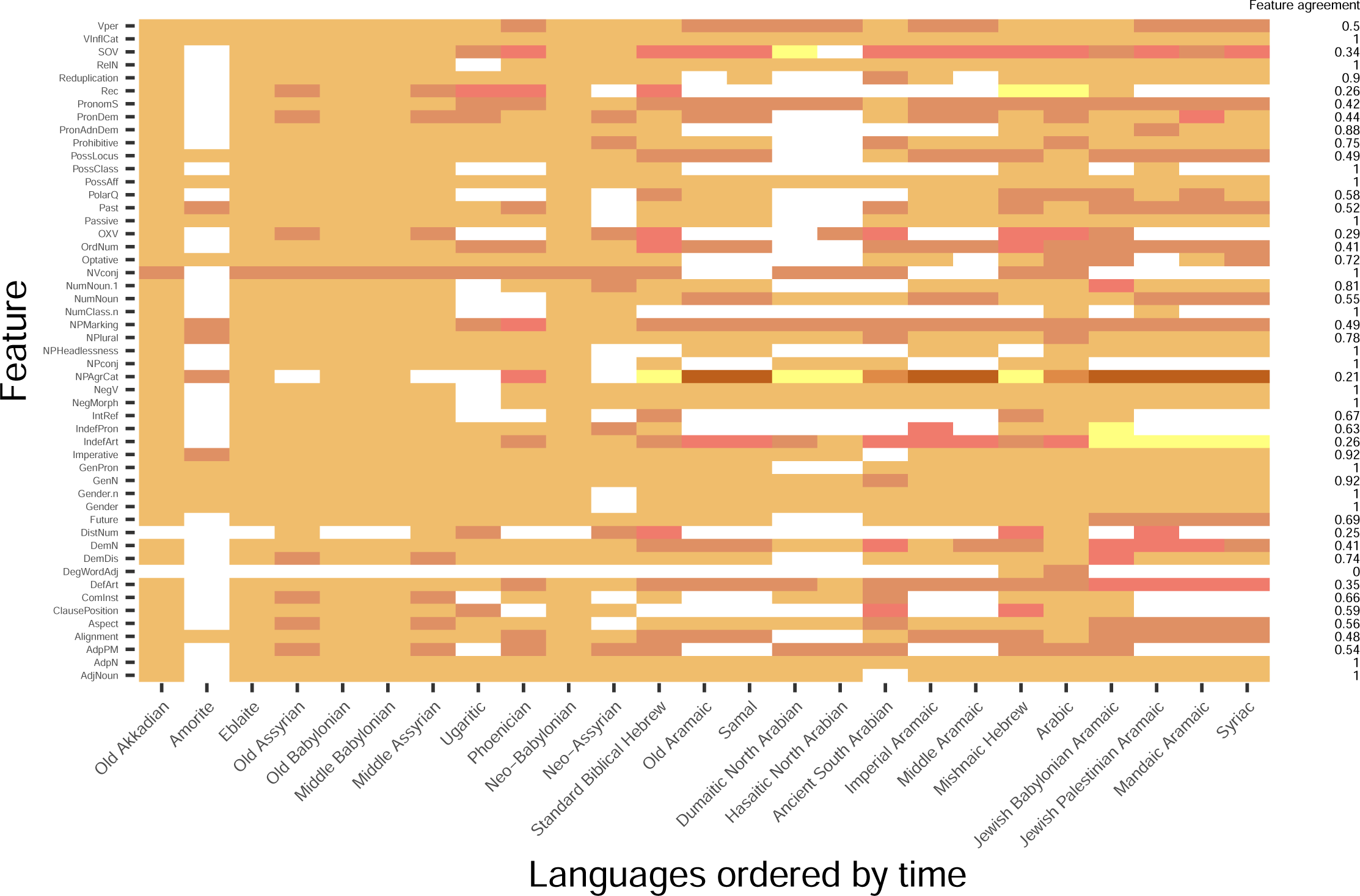
Differences in values for all the Semitic languages in all tested features. Different colors represent different values per feature. The values are different across rows. White stands for missing values. The rightmost column (*feature agreement*) shows the proportion of pairs of languages that share a feature (excluding missing values). The average agreement of feature values between language pairs is 0.69, with a standard deviation of 0.28 (see Supplementary Information Section S1.1 for the calculation of the feature agreement).

We then also manipulated the prior on universal preference, which effectively controls how much the feature distributions in a set of similar languages need to differ from the worldwide norm for sBayes to assign these languages to a cluster with a distinctive typological profile. A narrow prior on universal preference assumes that the worldwide feature distribution is known with certainty. In this case, already subtle deviations from the universal preference suffice for sBayes to report a deviating cluster. A weak prior assumes that the universal preference itself is uncertain. In this case, a feature is more readily explained by universal preference, attributing deviations to uncertainty rather than effects in a cluster. To assess the robustness of our results, we varied the precision *ρ* of the prior.

We find that the signal is stable for *ρ ∈ {*10, 30, 50, 70*}*, consistently reporting Hurrian-Sumerian as the only cluster not explained by universal or family-specific tendencies (Section S3.1.1, Supplementary Information). Only when the prior is very narrow (*ρ* = 90), sBayes also returns the remaining languages of the ANEA as deviating from the universal tendencies. This parallels the result of the regression analysis, which showed that the Ancient Near East languages are distinctive from universal tendencies only when focusing on a handful of feature states. Finally, we varied the number of possible clusters. We find decisive support for a model with a single cluster, suggesting that Hurrian-Sumerian is the only cluster deviating from the universal tendencies backed by the data (Section S3.1.3, Supplementary Information).

A possible confound of our results is that our data collapses different time periods. For example, Old Akkadian is attested around the mid-second millennium BCE, and Ugaritic is attested only a thousand years later in the late mid-first millennium BCE – but this difference does not inform our models. However, this is unlikely to have any effect on our findings. sBayes looks for similarities in the data, so subsetting the data would have made finding clusters less likely, not more, as there would be fewer languages per time slot. Our second finding, the Hurrian-Sumerian cluster, is also unlikely to have been caused by the conflation of time periods. First, (written) Sumerian and Hurrian do overlap in time for some 1,000 years, and second, other languages in the sample have the same time depth as Hurrian and Sumerian and were not assigned to this cluster (Egyptian, Akkadian, Eblaite, Elamite, Amorite, Hittite, Luwian, Phoenician, Ugaritic).

## 4 Discussion

Our analyses suggest that for the vast majority of features, the Ancient Near East largely follows the same universal profile as modern languages, suggesting a relatively stable distribution of linguistic features over time from antiquity to today. However, while the frequency distributions of the whole of the ANEA do not differ from current distributions, two languages from our sample clearly and robustly do: Hurrian and Sumerian. In what follows we first discuss the similarity between the ANEA and the rest of the world, and then assess the possible implications of the Hurrian-Sumerian cluster.

### 4.1 The ANEA as part of West Eurasia

All the languages (apart from Hurrian and Sumerian) in the sample fall within the posterior universal distributions from sBayes, consistent with the result from the regression analysis. This homogeneity is particularly interesting when considering the time depth of the sample. The data in the Ancient Near East sample ends in the mid-first millennium CE, while the data for the universal distribution is taken from modern languages, making the time distance between the two samples at least 1,500 years and at most over 5,000.

One interpretation of this finding is that the Ancient Near East was not very different from the modern distribution of linguistic features, supporting the notion of a time-independent and possibly stationary frequency distribution. However, the similarity between the ANEA and the rest of the world might be partly driven by the fact the ANEA has strong historical continuity with neighboring areas, leading to a linguistic profile that unites it with many present-day languages in our sample. There is in fact ample evidence for continued contact between the Ancient Near East through diplomatic, economic and technological relations extending all the way to the Indus Valley, Anatolia, Greece, the Caucasus, the Zagros mountains, North Africa, and south of the Nile river [36–38]. Naturally, the expansion of infrastructure, trade, and cultural exchange grew with time, reaching its peak towards the end of our sample, with the big empire conquests of the first millennia BCE and CE: the Persian Empire, the Hellenistic period, the Roman Empire, and the Islamic conquests, inevitably administering linguistic uniformity and wiping out endemic languages.

In addition to historical records and robust material evidence, linguistic evidence also tells a similar story of contact and spread of the languages of the Ancient Near East outwardly. For example, grammatical features (quantifiers, articles and other aspects of noun phrases) have spread throughout the Mediterranean, transforming the profiles of the languages there from earlier stages in an east-to-west trajectory of spread, making them similar to one another and discontinuous with their ancestor languages [39]. Some even suggest those spreads have reached as far north in Europe to reach the British Isles, where the Semitic languages are (controversially) argued to have left a traceable mark on the local Germanic and Celtic languages [40, 41]. One reconstruction of Proto-Indo-European proposes contacts with Semitic languages based on phonetics [42]. Though the overall theory is controversial, there are uncontested reconstructed lexical borrowings from Semitic into Proto-Indo-European (PIE), items such as ‘gold’ (PIE *\*Haus-*, Akkadian *hurāṣu*), ‘wine’ (PIE *\*woino-*, Semitic *\*wain-*) and ‘bull’ (PIE *\*tawro-*, Semitic *\*tawr-*), and these point to early trade and migration between the populations speaking those languages [43].

The relationship of contact and exchange between the Ancient Near East and neighboring areas is also evident from their demographic history, as reconstructed through population genetics. Genome-wide ancient DNA studies have revealed major differences between the genetic makeup of the different populations of the Near East at the time when agriculture entered the scene, around 12,000 YBP. At that time, the populations of Mesopotamia, the Levant and Anatolia were extremely genetically differentiated and little admixture between them is captured [44]. In the following millennia, agriculturalist populations from those regions have expanded in all directions, contributing to the genetic homogenization of West Eurasia [44, 45]. Genome-wide ancient DNA extracted from 22 individuals from Peqi’in cave in the southern Levant, dated to the fourth millennium BCE, shows that this population had received exogenous genetic influences from ancient Iranian and ancient Anatolian groups [46].

The connection between Ancient Near Eastern populations and European populations is also evident in the DNA of ancient European populations. Complete mitochondrial genomes of individuals from three different necropolises in Bulgaria dated to the third and second millennia BCE and associated with the Indo-European Thracian culture, include a mixture of lineages from East Europe and the Mediterranean, pointing to a mix of genetic sources as a result of population movements between the Mediterranean and Europe. This genetic influence from the Levant is not limited to Eurasia, as there is evidence for admixture west and southwards into Africa as well. The genomic makeup of Ethiopians is modeled to have a Eurasian component of a Levantine or Anatolian origin, dated to 3,000 years ago [44].

Thus, whereas the populations of the Near East before the Late Chalcolithic were distinct, the populations of the Levant coming from the Late Chalcolithic were already characterized by a degree of genetic homogeneity due to genetic admixture [46]. Languages start being recorded in writing not much after that, when our language sample begins. Hence, most of the languages in our sample typify multiple sources of ancestry that exhibit overall regional homogeneity due to sustained contact between speaker populations.

In summary, while the results from our regression analysis suggest that distributions might have been the same in the ANEA as now, we cannot rule out that this apparent homogeneity results from the specific historical continuity that the ANEA has had with neighboring regions.

### 4.2 Hurrian and Sumerian

The second finding of our study is the clustering of two languages that deviate from the homogeneity of the other languages of the ANEA: Hurrian and Sumerian. These two languages are two of the most ancient languages in the sample. Sumerian is the earliest known written language and as such the oldest language in our sample (Figure 1). While Sumerian has no known relatives (living or dead), Hurrian is part of the Hurro-Urartian language family [47]. Its only other known member is Urartian, the official language of Urartu in the ninth to sixth centuries BCE. Urartian is not part of our language sample due to lack of sufficient documentation. Even though Sumerian and Hurrian have been spoken in parallel for some centuries, their geographical distance is considerable. Sumerian was spoken in southern Mesopotamia, and Hurrian was spoken in Anatolia and northern Mesopotamia. Though both languages were in contact with the oldest known Semitic language, Akkadian, there is little evidence of direct contact between Sumerian and Hurrian. This makes it very unlikely that the Sumerian-Hurrian cluster reflects a distinct, narrow linguistic area of its own.

We do not know what the linguistic landscape around them was like. We have no lingering traces of the contemporaneous languages which were not committed to writing, and no insight as to which languages were spoken before writing appeared on the scene altogether and their possible typology. Therefore, two possible explanations of the exceptional status of the two languages arise. Either Hurrian and Sumerian are the last survivors of an earlier regional linguistic area of unknown extent, which was wiped out by the spread of the Semitic and Indo-European languages, or even more broadly, Hurrian and Sumerian reflect an ancient global distribution different from today’s.

Under either interpretation, the data from Hurrian and Sumerian suggest a massive transformation of the linguistic landscape in the last few millennia. This finding is consistent with the fact that many other lineages in the Near East, beyond those of Hurrian and Sumerian, such as Elamite and Egyptian in our sample, along with whole branches and families we have no remnants of, were gradually replaced by Semitic and Indo-European languages, further changing the linguistic distributions in the region. The phenomenon of language replacement and typological flattening is not restricted to the Ancient Near East. Well-documented cases are the Indo-European spread in west Eurasia [13] and the Bantu expansion in Sub-Saharan Africa [14].

We cannot presently distinguish the two scenarios, but there is tentative evidence to support the second. Some of the features that differentiate Hurrian and Sumerian from the rest of the sample (such as alignment, verbal inflectional categories, possessive classes, and optative, Section S3.2.4, Supplementary Information) are also shared by languages that are found in areas of the world referred to in the literature as “enclaves” or “residual zones” in Eurasia, i.e. regions that have been postulated to retain the linguistic characteristics of Eurasia before the major spread of modern language families [17, 48, 49].

Indeed, the distributions of some structural features tentatively suggest a possible link between the frequency distributions of Hurrian and Sumerian and the current-day residual zones such as the Caucasus and the Himalayas, although much richer data would be needed to test these links formally. Candidate features are morphological optatives, the number of categories in maximally inflected verb forms (sometimes called ‘degree of synthesis’), ergative alignment, and genderless personal pronouns.

Morphological optative refers to the presence of a verbal inflectional category dedicated to the expression of the speaker’s wish or desire [50, 51]. It does not refer to other grammatical means of marking the speaker’s wish which are not inflectional, such as a modal verb (English *may*). Both Sumerian and Hurrian had an inflectional optative. Morphological optative is rare in Eurasia today, but is still present in enclave languages (Figure 6a). It has been a common feature in the languages of the past and disappeared with time. In addition to Hurrian and Sumerian, morphological optatives are also abundant in the rest of the ancient Near Eastern languages such as Egyptian, Akkadian, Hebrew and Ugaritic. Many ancient languages of the Indo-European family such as Ancient Greek [52], Old Iranian [53, Old Persian and Avestan], and Vedic Sanskrit [54] also have a morphological optative. Most modern successors of the ancient languages with inflectional optative lost it, and there is reason to believe that inflectional optative may have been common in the past, reinforced by biases in language families that have died off since, and by language contact. Since the extinction of major lineages carrying that feature, the presence of it in the region diminished. In the Caucasus, though, areal pressure seems to be influential in maintaining this feature.

**Figure 6:**
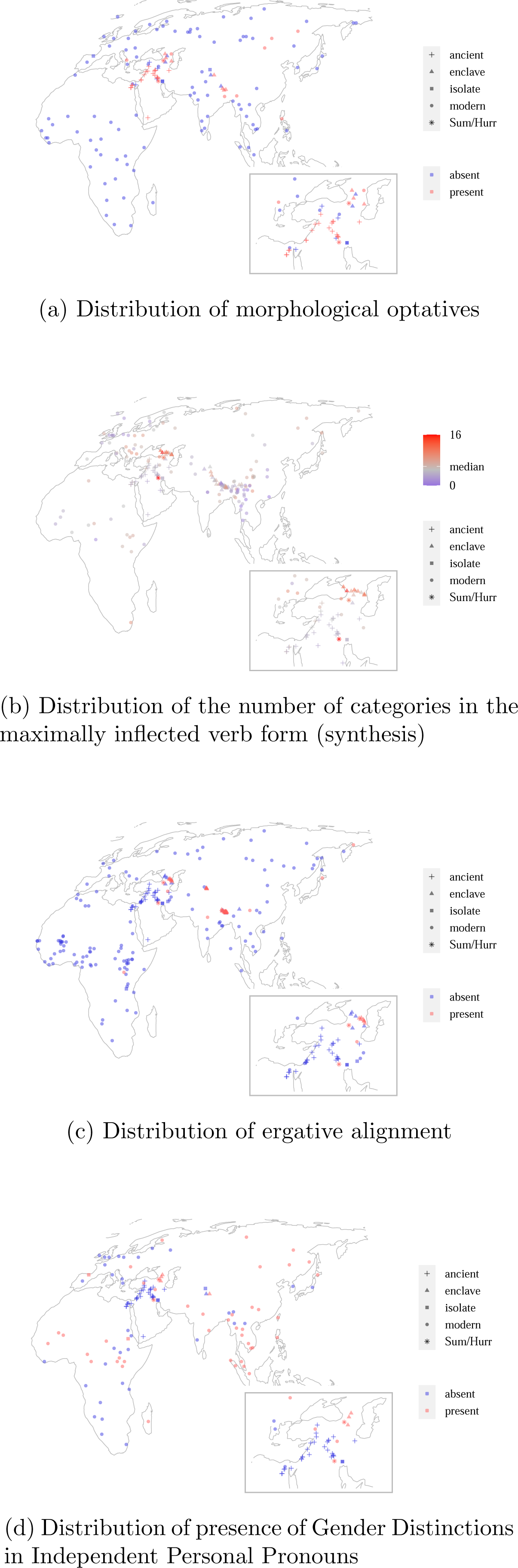
Distribution of features that may have been more prominent in prehistory. Insets of the Near East common in the past, but have diminished with time. The diminished degree of synthesis could possibly be attributed to universal pressures disfavoring very high verbal synthesis (for example, cognitive pressures), making it easily lost through time or areal diffusion.

Inflectional synthesis refers to the use of verbal inflection to express grammatical relations [48]. In other words, when a grammatical category is expressed through bound inflection, it is a synthetic rather than analytic realization of that category. Today, Eurasia is marked by relatively low synthesis of the verb. Exceptions to that are the Himalayas and the Caucasus, where a high degree of synthesis is clustered (Figure 6b). As a high degree of synthesis is shared by Sumerian and Hurrian and the enclave languages, higher levels of synthesis may have been

Morphosyntactic alignment describes the type of grammatical relationship between the arguments of the verb. Arguments are aligned when they are marked in the same way, under a certain condition [55, 56]. In an ergative alignment, the sole (S) argument of an intransitive verb (e.g. ‘work’ or ‘sleep’) is marked in the same way as the patient (P) and differently from the agent (A) argument of a transitive verb (e.g. ‘build’ or ‘hit’): (*S* = *P ̸*= *A*). Though not much is known about Hurrian’s sister language, Urartian, the available inscriptions also exhibit an ergative alignment [57], suggesting strongly that this was a family-wide trait. Of all ancient Indo-European languages Hittite is the only one with ergative alignment. The development of ergativity in Hittite perhaps indicates an areal pressure in ancient times [58].

Lastly, independent personal pronouns are lexical items capable of carrying stress, that are not bound forms (such as the bound person affixes and clitics in previous examples) [59]. A language may distinguish gender in its independent personal pronouns. Both Hurrian and Sumerian lack any gender distinction in independent personal pronouns. Unlike the other features surveyed here, genderless pronouns are the most common pattern in the world today (Figure 6d). They are also common in the enclaves and isolates. Gendered pronouns are common in Africa in the Niger-Congo and Afro-Asiatic languages, and in the Indo-European languages of Europe [59]. These three language families are associated with the agricultural spreads of the Neolithic, pointing to an ancient pattern of genderless pronouns, as seen in Sumerian and Hurrian and kept in the enclaves and isolates today. Intriguingly, latecomers into the enclaves, such as Ossetic in the Caucasus and Nepali in the Himalayas (both Indo-European), adapted to the areal pattern and lost their gender distinction in independent personal pronouns, promoting further the hypothesis that this feature is subject to strong areal pressures.

To sum up, Hurrian and Sumerian stand out from both the ANEA and the global sample, suggesting a formerly unknown ancient regional or global distribution.

## 5 Conclusions

Together, our findings challenge the notion that present-day frequency distributions of linguistic features are representative of those distributions in earlier times, or even more so, are representative independently of any given time. The global distribution of linguistic features is therefore not exempt from radical change and hence not necessarily representative of the language faculty, but may be contingent on historical events and cultural evolution.

## Acknowledgment

We would like to thank Dr. Chiara Barbieri for her consultation on the interpretation of the genetic data presented in this paper. We would like to thank the University Research Priority Program (URPP) Language and Space, University of Zurich, Zurich, Switzerland, for supporting this project.

## Footnotes

1 Hurro-Urartian is a language family comprising of two known languages: Hurrian and Urartian. There are not enough surviving documents of Urartian for constructing its full grammar and, therefore, it is not represented in our dataset and Hurrian is considered effectively a language isolate. Language isolates are languages which are the sole survivors of their families, rather than languages that have developed in isolation

