## Supplementary Information for "Oldest attested languages in the Near East reveal deep transformations in the distribution of linguistic features"

This PDF file includes:
Supplementary Text
Figs. S1 to S12
Table S1

#### S1 Language and feature selection and recoding

The selection of language varieties was done based on data availability and documentation. Data was collected for every doculect (documented language variety) from the region. Therefore, some languages in the sample are represented by a few language varieties. For example, Aramaic, which was well documented, is represented by eight distinct varieties: Old Aramaic, Samal, Imperial Aramaic, Middle Aramaic, Jewish Palestinian Aramaic, Jewish Babylonian Aramaic, Syriac, Mandaic Aramaic. Some are different diachronic stages of the languages, whereas some are different geographic dialects. Other languages, though having many doculects, are represented by one language variety in the sample. Different doculects are combined to one representative variety in the case where all doculects do not exhibit any distinction in the morphosyntactic features that were collected. An example for such case is Arabic. Morphosyntactic data was collected for 10 doculects: Classic Standard Arabic, pre-Islamic Hijazi, ˀAzdi, Yemeni, Southern Yemeni, Huḏayli, Ṭayyiˀi, Qaysi, ˀAsadi, and Tamīmi. But due to either too sparse attestation, inability to distinguish a morphological difference because of the writing system (vowel quality for example), or actual morphologicl similarity, all 10 doculects were identical with regards to the morphosyntactic features we looked at, and hence are represented by one variety 'Arabic'.

The selection of linguistic features was guided by the motivation to achieve as much descriptive comparability between the languages in our samples as possible. For this reason, we used well-established morphological features in our analysis, described and collected for a vast number of the world's languages in two large databases: WALS [1] and AUTOTYP [2]. We selected all 51 relevant grammatical features from both databases (Table S1), excluding duplicates appearing in both, even if coded differently, keeping only one.

In the regression analysis, categories are assumed to be at equal distance, which is not the case for features where one language can have more than one category (e.g., *mixed* or *both*). In this case, the distance between categories *A* and *B* is different from the distance between categories *A* and *A&B*. In response to this we adapted the WALS data transformations set-up by [3] (`https://github.com/IVS-UZH/WALS-recodings`), and binarized features with *mixed*/*both* categories into the two features 'category A presence' and 'category B presence', both with categories *true* and *false*. Languages with *mixed*/*both* as a value for the original feature now have a *true* value for both 'category A presence' and 'category B presence'.

This systematic recoding is only feasible for otherwise binary features. For features with more categories, we cannot generally know which ones contributed to the *mixed* value of the language. For example, for the category *no dominant order* in the feature 'Word Order SOV', wich has six possible categories, we cannot reconstruct which of the six possible word orders are present in the language in such a case. For those features, we excluded the category *mixed* from the analysis and, where possible, filled in the missing data. For the recoded AUTOTYP features, the data contained the categories that contributed to the *mixed* category, making it possible to retain all information. These features were recoded by hand because of inconsistencies in the coding. In total, we recoded nine of the features containing a *mixed/both* category, yielding a set of 70 morphological features.

#### S1.1 List of features and feature agreement

*Original feature name* is the name under which the feature appears in the WALS or AUTOTYP databases; *Recoded feature name* is the name the feature has after running it through the recoding code; and *Abbreviated feature name* is what is the abbriviated version under which the features can be found in our analysis.

**Feature agreement.** In Figure 5 of the main article we show the agreement of feature values between language pairs as an indicater of a feature's stability. This agreement value is calculated in the following way. For each feature $f$ with $N_f$ possible states $S_f = \{1, ..., N_f\}$, we denote the number of occurences of each state $s \in S_f$ in our dataset by $k_s$. $K_f = \sum_{s \in S_f} k_s$ denotes the total number of values for this feature. Note that missing values are not counted in $k_s$ or $K_f$. For each state $s$ there are $\binom{k_s}{2}$ pairs of languages that share this feature state. In total there are $\binom{K_f}{2}$ language pairs that may or may not share a state. Consequently, the fraction of language pairs that share a feature state in feature $f$ is:

$$\text{agreement}(f) = \frac{\sum_{s \in S_f} \binom{k_s}{2}}{\binom{K_f}{2}}$$

| Original feature name | Recoded feature name | Abbreviated feature name |
|---|---|---|
| Reciprocal Constructions | Reciprocal.Presence | Rec.P |
| Reciprocal Constructions | ReciprocalReflexive.Identical | RecRef.I |
| Reciprocal Constructions | ReciprocalReflexive.Distinct | RecRef.D |
| Position of Interrogative Phrases in Content Questions | ContentQInitial | ContQInitial |
| Position of Interrogative Phrases in Content Questions | ContentQNonInitial | ContQNonInitial |
| Order of Numeral and Noun | NumNoun | NumNoun |
| Order of Numeral and Noun | NounNum | NounNum |
| Order of Degree Word and Adjective | DegWordAdj | DegWAdj |
| Order of Degree Word and Adjective | AdjDegWord | AdjDegW |
| Order of Genitive and Noun | GenN | GenN |
| Order of Genitive and Noun | NGen | NGen |
| Order of Adjective and Noun | AdjNoun | AdjN |
| Order of Adjective and Noun | NounAdj | NAdj |
| Position of Pronominal Possessive Affixes | PossAffixes.Presence | PossAff.P |
| Position of Pronominal Possessive Affixes | PossPrefix | PossPre |
| Position of Pronominal Possessive Affixes | PossSuffix | PossSuff |
| Comitatives and Instrumentals | | ComInst |
| Noun Phrase Conjunction | | NPconj |
| Nominal and Verbal Conjunction | | NVconj |
| Definite Articles | | DefArt |
| Indefinite Articles | | IndefArt |
| Expression of Pronominal Subjects | | PronomS |
| Indefinite Pronouns | | IndefPron |
| Third Person Pronouns and Demonstratives | | PronDem |
| Distance Contrasts in Demonstratives | | DemDis |
| Pronominal and Adnominal Demonstratives | | PronAdnDem |
| Intensifiers and Reflexive Pronouns | | IntRef |
| Gender Distinctions in Independent Personal Pronouns | | GenPron |
| Systems of Gender Assignment | | Gender |
| Number of Genders | | Gender.n |
| Negative Morphemes | | NegMorph |
| Coding of Nominal Plurality | | NPlural |
| Ordinal Numerals | | OrdNum |
| Distributive Numerals | | DistNum |
| Order of Adposition and Noun Phrase | | AdpN |
| Order of Demonstrative and Noun | | DemN |
| Order of Negative Morpheme and Verb | | NegV |
| Order of Object Oblique and Verb | | OXV |
| Order of Relative Clause and Noun | | RelN |
| Passive Constructions | | Passive |
| The Optative | | Optative |
| The Prohibitive | | Prohibitive |
| The Morphological Imperative | | Imperative |
| Person Marking on Adpositions | | AdpPM |
| Position of Polar Question Particles | | PolarQ |
| Possessive Classification | | PossClass |
| Locus of Marking in Possessive Noun Phrases | | PossLocus |
| Reduplication | | Reduplication |
| The Future Tense | | Future |
| The Past Tense | | Past |
| Perfective Imperfective Aspect | | Aspect |
| Verbal Person Marking | | Vper |
| Order of Subject Object and Verb | | SOV |
| NPMarking | Head.driven.agr | Head.driven.agr |
| NPMarking | Construct.state | Cons.state |
| NPMarking | Juxtaposition | Juxtaposition |
| NPMarking | Governed | Governed |
| NPAgrCat | Number | Agr.Number |
| NPAgrCat | Gender | Agr.Gender |
| NPAgrCat | Case | Agr.Case |
| NPAgrCat | class | Agr.class |
| NPAgrCat | mixed | Agr.mixed |
| NPAgrCat | Definiteness | Agr.Def |
| NPAgrCat | State | Agr.State |
| NPAgrCat | none | Agr.none |
| NPHeadlessness | | NPHeadlessness |
| ClausePosition | | ClausePosition |
| NumClass.n | | NumClass.n |
| AlignmentCaseDominantSAP | | Alignment |
| VInflCatandAgrFmtvMax.binned3 | | VInflCat |

Table S1: Feature list

#### S1.2 List of languages

| Branch | Language | Language Variety | Time | Source |
|---|---|---|---|---|
| Egyptian | Egyptian | Earlier Egyptian | 3,200 BCE-2,000 BCE | [4], [5], [6] |
| | Egyptian | Middle Egyptian | 2,000 BCE-1,300 BCE | [4], [5], [6] |
| | Egyptian | Later Egyptian | 1,350 BCE-300 CE | [4], [5], [6] |
| | Coptic | Coptic | 200 CE-1,500 CE | [7] |
| East Semitic | Eblaite | Eblaite | 2,300 BCE-2,250 BCE | [8], [9], [10] |
| | Akkadian | Old Akkadian | 2,500 BCE-1,950 BCE | [8], [9], [10] |
| | Akkadian | Old Assyrian | 1,950 BCE-1,750 BCE | [11] |
| | Akkadian | Middle Assyrian | 1,500 BCE-1,000 BCE | [11] |
| | Akkadian | Neo-Assyrian | 1,000 BCE-600 BCE | [12], [11] |
| | Akkadian | Old Babylonian | 1,950 BCE-1,530 BCE | [8], [9], [10] |
| | Akkadian | Middle Babylonian | 1,530 BCE-1,000 BCE | [8], [9], [10] |
| | Akkadian | Neo-Babylonian | 1,000 BCE-625 BCE | [8], [9], [10] |
| Northwest Semitic | Amorite | Amorite | 2,500 BCE-1,200 BCE | [13] |
| | Phoenician | Phoenician and Punic | 1,200 BCE-600 BCE | [14], [15], [16] |
| | Ugaritic | Ugaritic | 1,300 BCE-1,190 BCE | [17], [18] |
| | Hebrew | Standard Biblical Hebrew | 1,000 BCE-100 CE | [19], [20], [21, 22] |
| | Hebrew | Mishnaic Hebrew | 100 CE-500 CE | [19], [20], [21], [22, 23] |
| | Aramaic | Old Aramaic | 900 BCE-600 BCE | [24], [25] |
| | Aramaic | Samal | 800 BCE-700 BCE | [24], [26] |
| | Aramaic | Imperial Aramaic | 600 BCE-200 BCE | [24], [27] |
| | Aramaic | Middle Aramaic | 200 BCE-250 CE | [24] |
| | Aramaic | Jewish Palestinian Aramaic | 200 CE-1,200 CE | [24], [28], [29] |
| | Aramaic | Jewish Babylonian Aramaic | 200 CE-1,200 CE | [24], [30] |
| | Aramaic | Syrian | 200 CE-1,200 CE | [24], [31], [32] |
| | Aramaic | Mandaic | 200 CE-1,200 CE | [24], [33] |
| Central Semitic | Epigraphic South Arabian | Epigraphic South Arabian | 700 BCE-600 CE | [34], [35] |

| | | | | |
|---|---|---|---|---|
| | Ancient North Arabian | Dumaitic | 800 BCE-400 CE | [36], [37] |
| | Ancient North Arabian | Hasaitic | 800 BCE-400 CE | [36], [37] |
| | Arabic | Arabic | 200 CE-700 CE | [38], [39] |
| Anatolian | Hittite | Hittite | 1,650 BCE-1,190 BCE | [40] |
| Anatolian | Luwian | Luwian | 1,140 BCE-700 BCE | [41] |
| Isolate | Hurrian | Hurrian | 2,300 BCE-1,000 BCE | [42] |
| | Sumerian | Sumerian | 3,350 BCE-1,800 BCE | [43] |
| | Elamite | Old Elamite | 2,600 BCE-550 BCE | [44], [45] |
| | Elamite | Achaemenid Elamite | 550 BCE-330 BCE | [44], [45] |

Table S2: List of languages and sources

*Branch* is the higher level phylogenetic branch the language belongs to; *Language* is the name of the language the language variety belongs to; *Language variety* is the local/diachronic lect described in the data; *Time* indicates the duration of attestation of the language variety; and *Source* include the grammars used to gather the grammatical data.

#### S2 Regression results

The regression coefficients show the influence of the area (i.e. being within or outside the ANEA) on the probability of each state to be present. A positive coefficient means that the variable state in question is more likely in the Ancient Near East than elsewhere. A negative coefficient indicates that it is more likely outside the Ancient Near East. Figures S1 and S2 show $50\%$ and $90\%$ credible intervals for the coefficients of each state based on highest posterior probability density estimates. Credible intervals centered around $0$ indicate little or no areal signal. In the figures, the labels on the y-axis give the abbreviated feature names (see Table S1), followed by the state for which the regression coefficient was estimated. For example, *IndefPron Spe* estimates the effect of area on the presence of the state *Spe* of the feature *IndefPron*.

Bayesian regression requires us to define a prior distribution on the regression coefficients. We use a normal distribution $N(0, \sigma^2)$ as a prior. We set the standard deviation to $\sigma = 1$ [46].

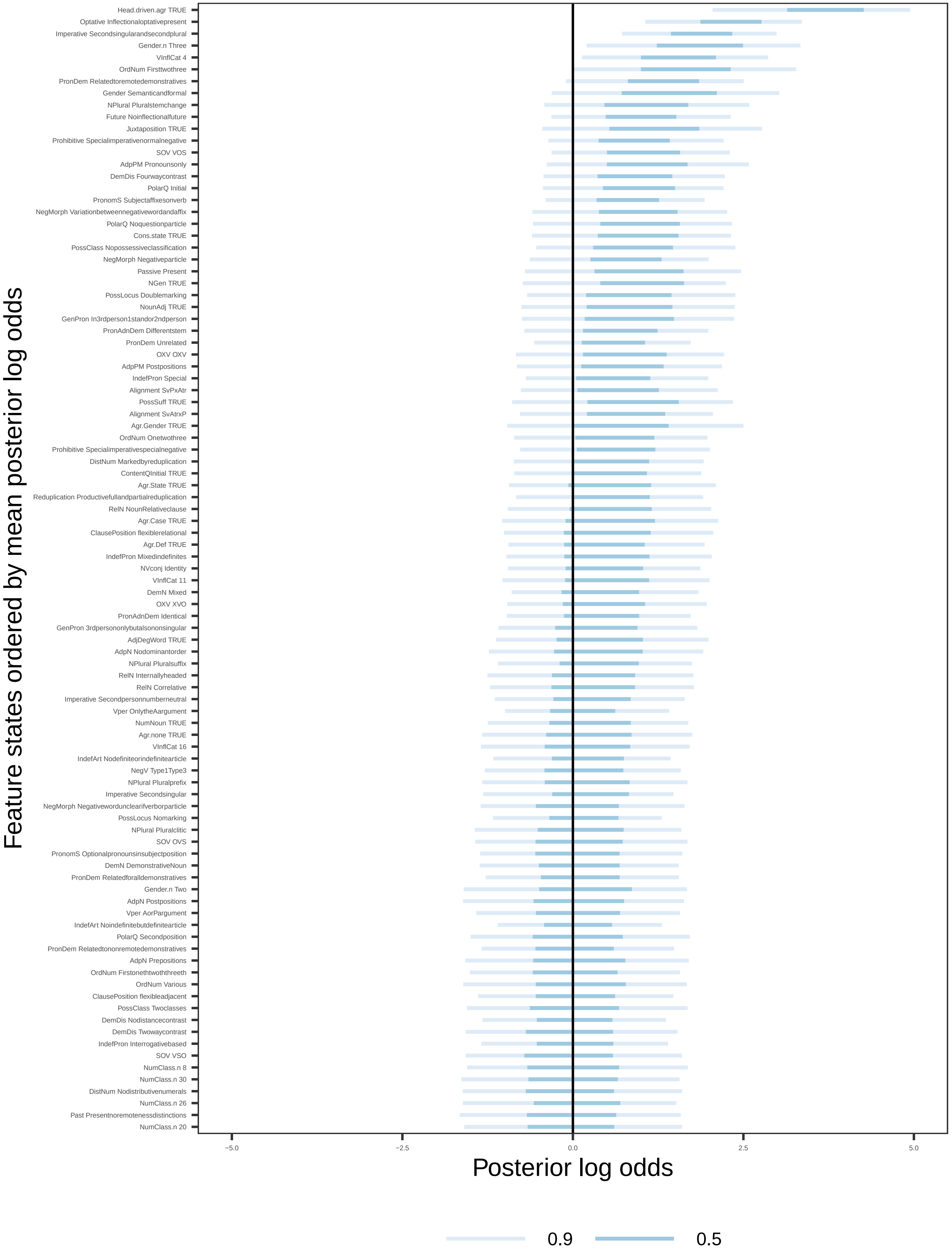


Figure S1: The effect of area (within the Ancient Near East vs. outside of it) on the likelihoods of grammatical traits with feature-state labels. (1/2)

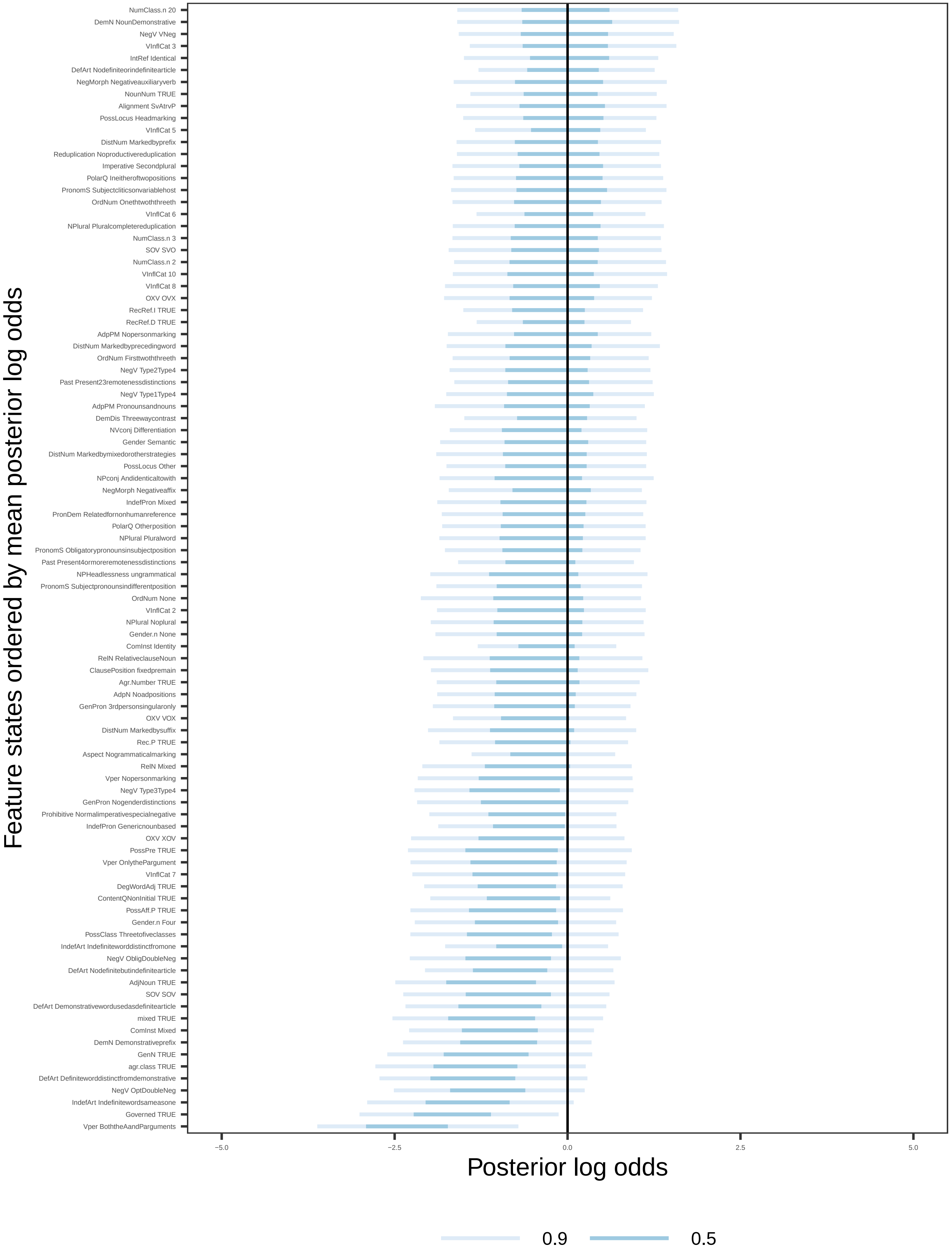


Figure S2: The effect of area (within the Ancient Near East vs. outside of it) on the likelihoods of grammatical traits with feature-state labels. (2/2)

### S3 sBayes results

#### S3.1 Sensitivity analyses

We performed a series of sensitivity analyses to verify that the sBayes results are stable for varying parameter configurations. Specifically, we varied the prior for universal preference, turned the mixture component for inheritance on and off and gradually increased the number of areas.

##### S3.1.1 Varying universal prior precision ($\rho$)

The universal distribution defines the probability over feature states not explained by inheritance or assigned to a cluster. The universal distribution is parameterized by $\alpha_{f,s}$, the probability of observing state $s$ in feature $f$. We use a Dirichlet prior on $\alpha$, with mean $\mu$ and precision or strength $\rho$. We inform the the prior with a global stratified sample of 100 languages, representing each feature's worldwide norm. The mean of the worldwide norm directly defines the mean of the prior. We treat the precision as a hyperparameter and vary its value in iterative runs.

A low precision results in a wide prior, implying that the worldwide norm is uncertain. In that case, the universal distribution is estimated locally from the data at hand with little influence from the stratified sample. The languages emerging as an area differ from the local background, but less so from the worldwide norm. A high precision results in a narrow prior, implying that the worldwide norm is known with certainty. In this case, the stratified sample strongly influences universal preference estimates. The languages in the area differ from the worldwide norm, but less so from the local background distribution. The stronger the prior, the more confident the model is about the worldwide norm. In this case, already subtle differences in a set of languages suffice for sBayes to assign them to an area, assuming, of course, that the languages are similar.

We carry out sensitivity analyses where we gradually increase the precsision between $10$ and $90$ to explore the influence of the hyperparameter on the inferred areas. For each experiment, we visualise the most salient areal signal in a consensus map. The consensus map includes only those languages that are together in a cluster in a minimum proportion of posterior samples, e.g., more than 20% (thin grey line in Figure S3).

Figure S3 shows the consensus maps for $\rho \in \{10, 30, 50, 70, 90\}$. The results consistently confirm the Hurrian-Sumerian link for $\rho \in \{10, 30, 50, 70\}$. Only for a very narrow and, thus, strict prior on the universal distribution, sBayes estimates a large area comprising most of the languages in our sample.

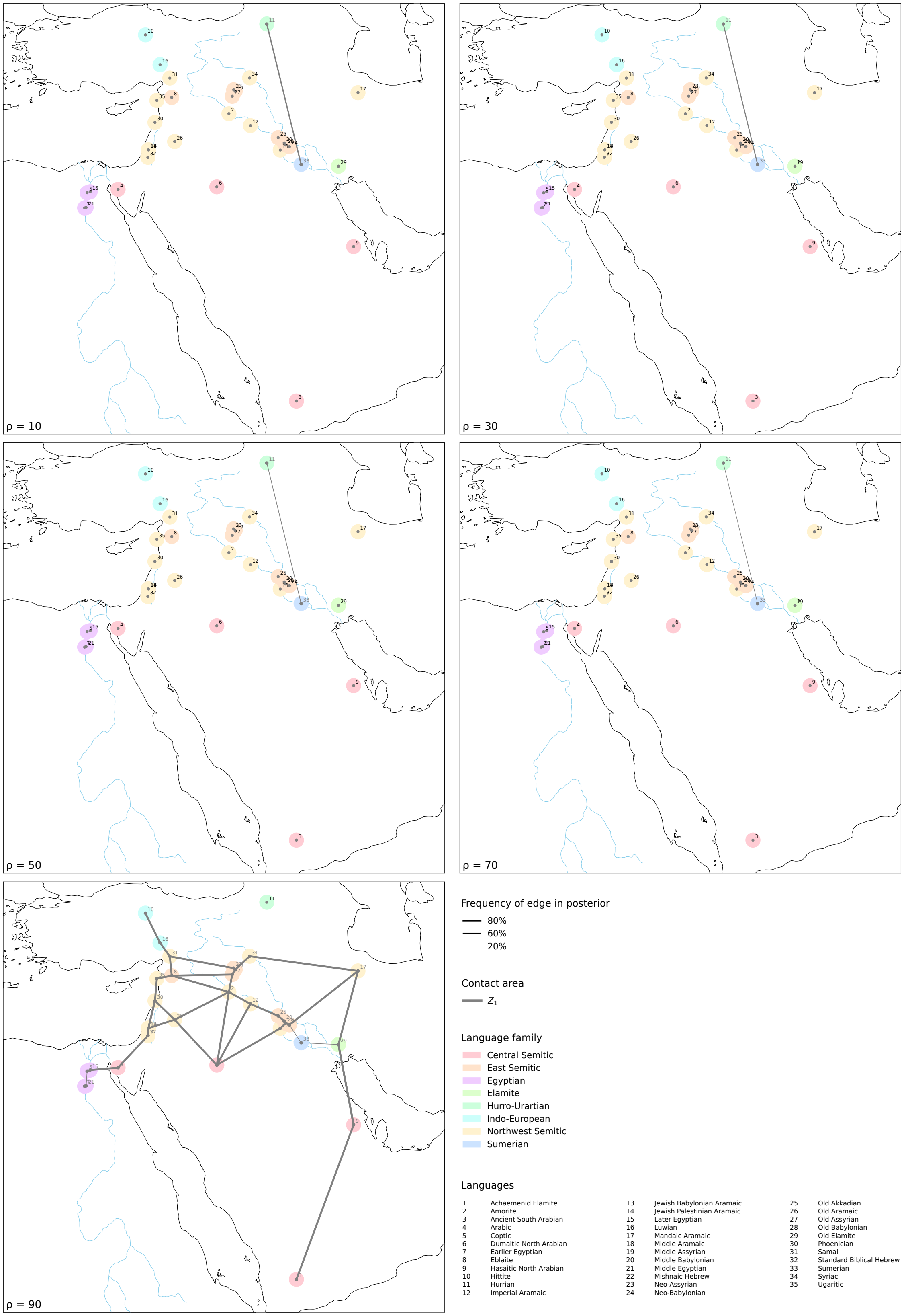


Figure S3: Maps of the consensus area for $\rho \in \{10, 30, 50, 70, 90\}$.

##### S3.1.2 Turning off the mixture component for inheritance

By turning of the inheritance component (setting its weight to 0), we exclude all information on phylogenetic relatedess from the analysis and force sBayes to allocate all evidence to either universal preference or a cluster. The results are the same as for models that include inheritance (Figure S4).

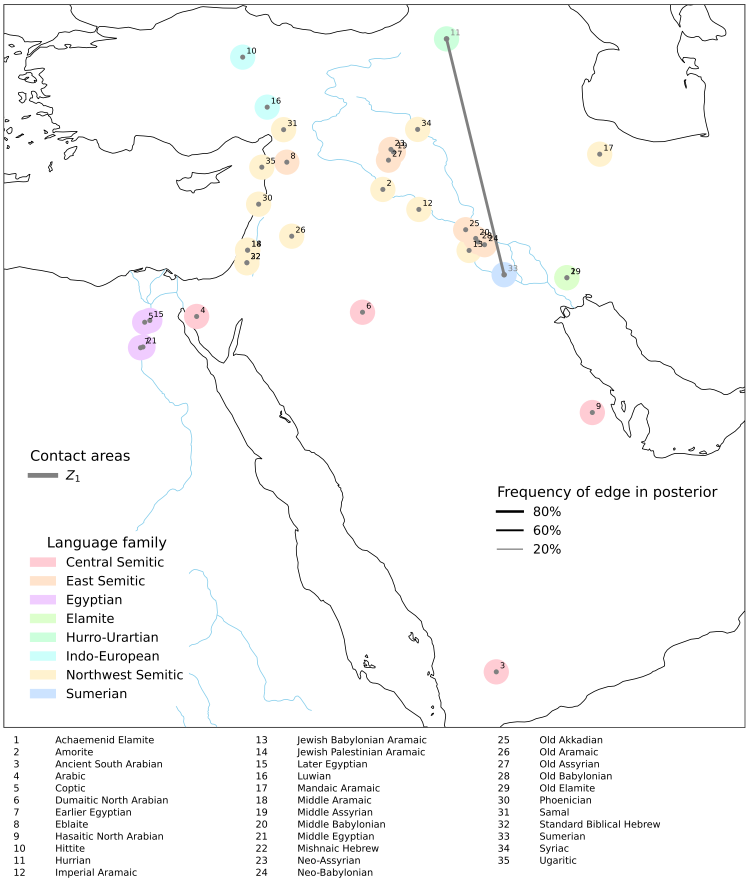


Figure S4: Posterior consensus map for an sBayes analysis without a mixture component for inheritance.

###### S3.1.3 Varying number of areas

We increased the number of areas, $K$ and compared the support for each model with the deviance information criterion (DIC) (see, ranacher2021sbayes). The inclusion of additional areas increases the DIC, e.g. by over 600 logarithmic units from $K = 1$ to $K = 2$, indicating decisive support for a single area.

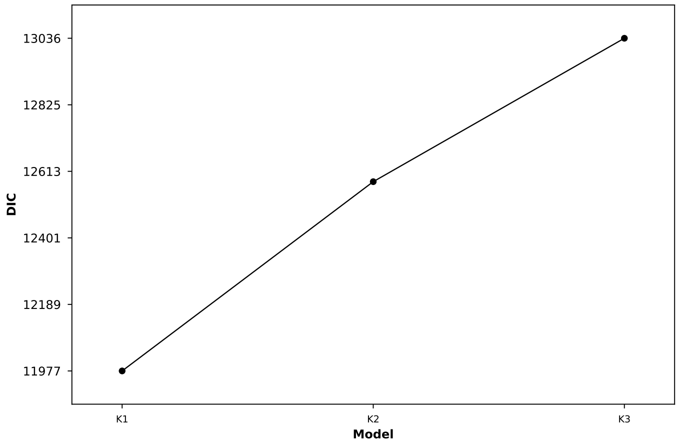


Figure S5: The deviance information crtierion for an increasing numbe rof areas $K$.

##### S3.2 Inferred parameters

We report all relevant inferred parameters for the main analysis ($\rho = 10$). Specifically, we plot the inferred assignment of languages to areas, and per feature, the universal preference ($\alpha$) the preference in the area ($\gamma$) and the weights of each mixture component. For $alpha$ and $gamma$, we always show a kernel density estimate (KDE) of the posterior state distribution. The shape of the plot for $\alpha$ and $\gamma$ depends on the number of states in a feature. For features with states, the $x$-axis shows the value for the state preference, and the $y$-axis shows the KDE of the posterior distribution. For features with more states, we plot a simplex with one corner per state. The distance to the corners indicates the preference of a state. The inferred posterior distribution over the state preferences is shown as a heat map in the simplex, with darker colours indicating higher densities and lighter colors indicating lower densities.

###### S3.2.1 Assignment of languages to areas

For each language $l$, `sBayes` estimates the probability that $l$ is part of the area $Z_1$. Here, we plot this posterior probability for each language in a pie chart.

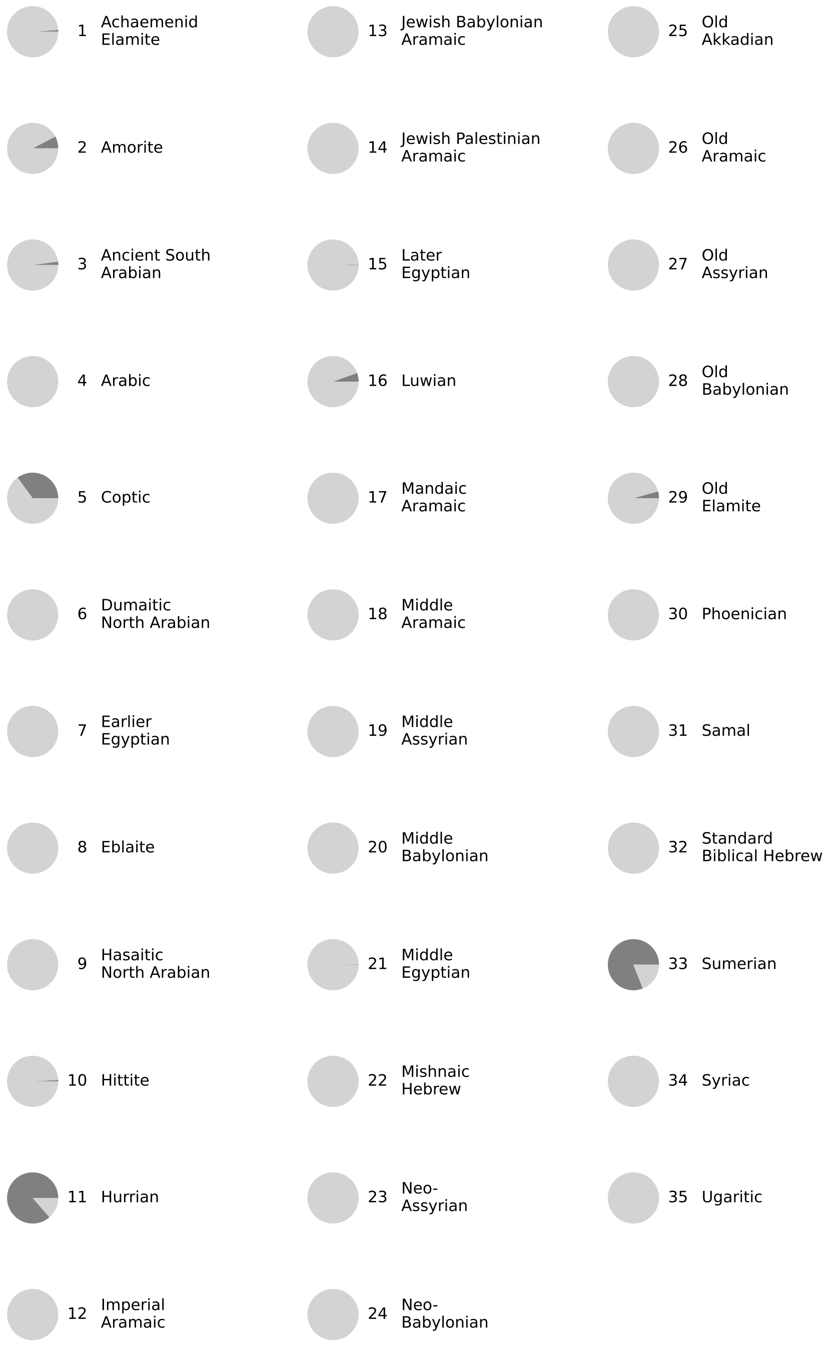


Figure S6: The assignment of languages to the area $Z_1$. The circle represents all posterior samples. The dark grey segment corresponds to the fraction of posterior samples where the language is assigned to $Z_1$.

##### S3.2.2 Universal preference

The following pages show state state probabilities, $\alpha$, for each feature $f$ in the universal distribution.

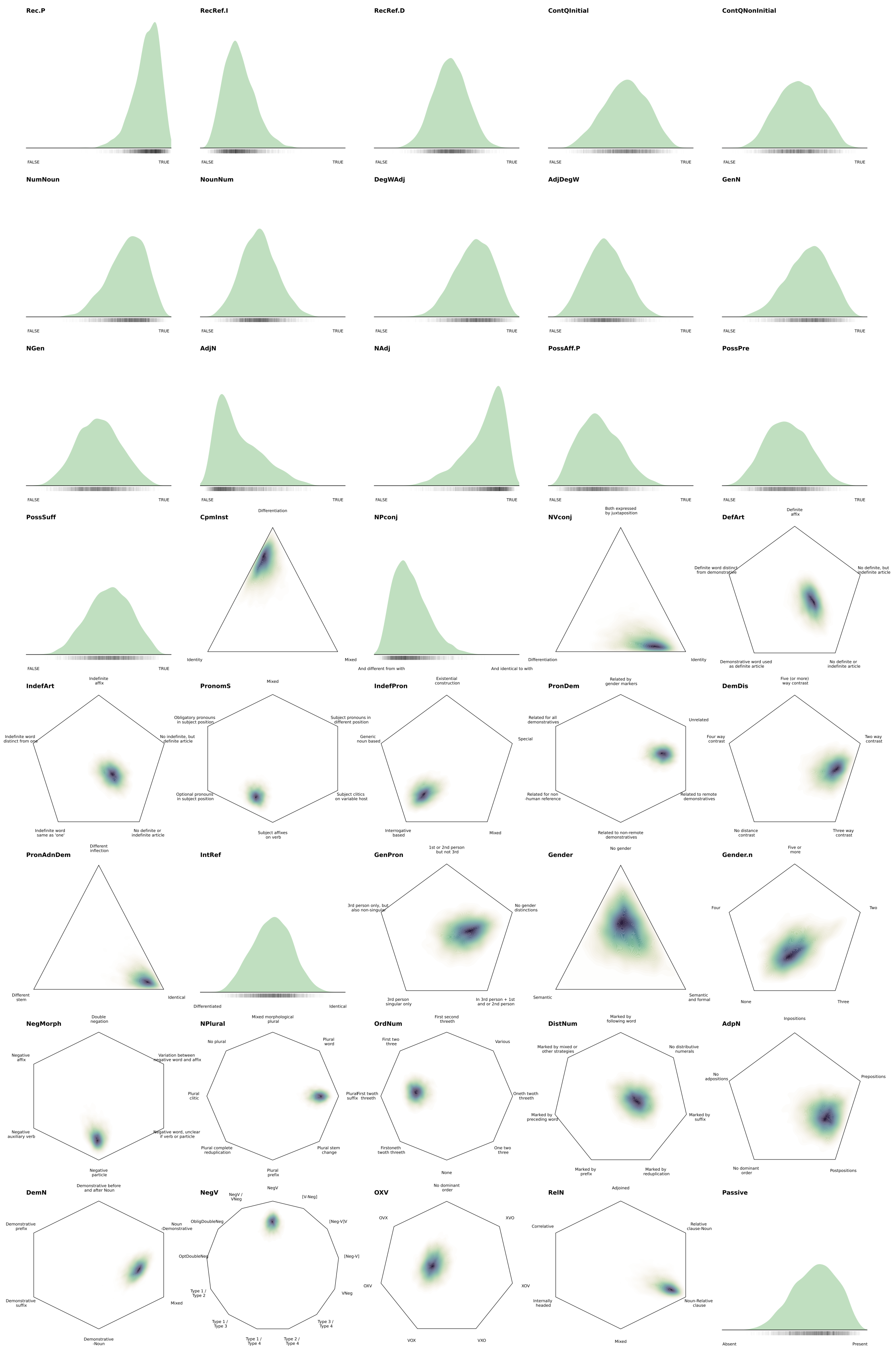


Figure S7: The state probabilities $\alpha$ for each feature in the universal distribution. (1/2)

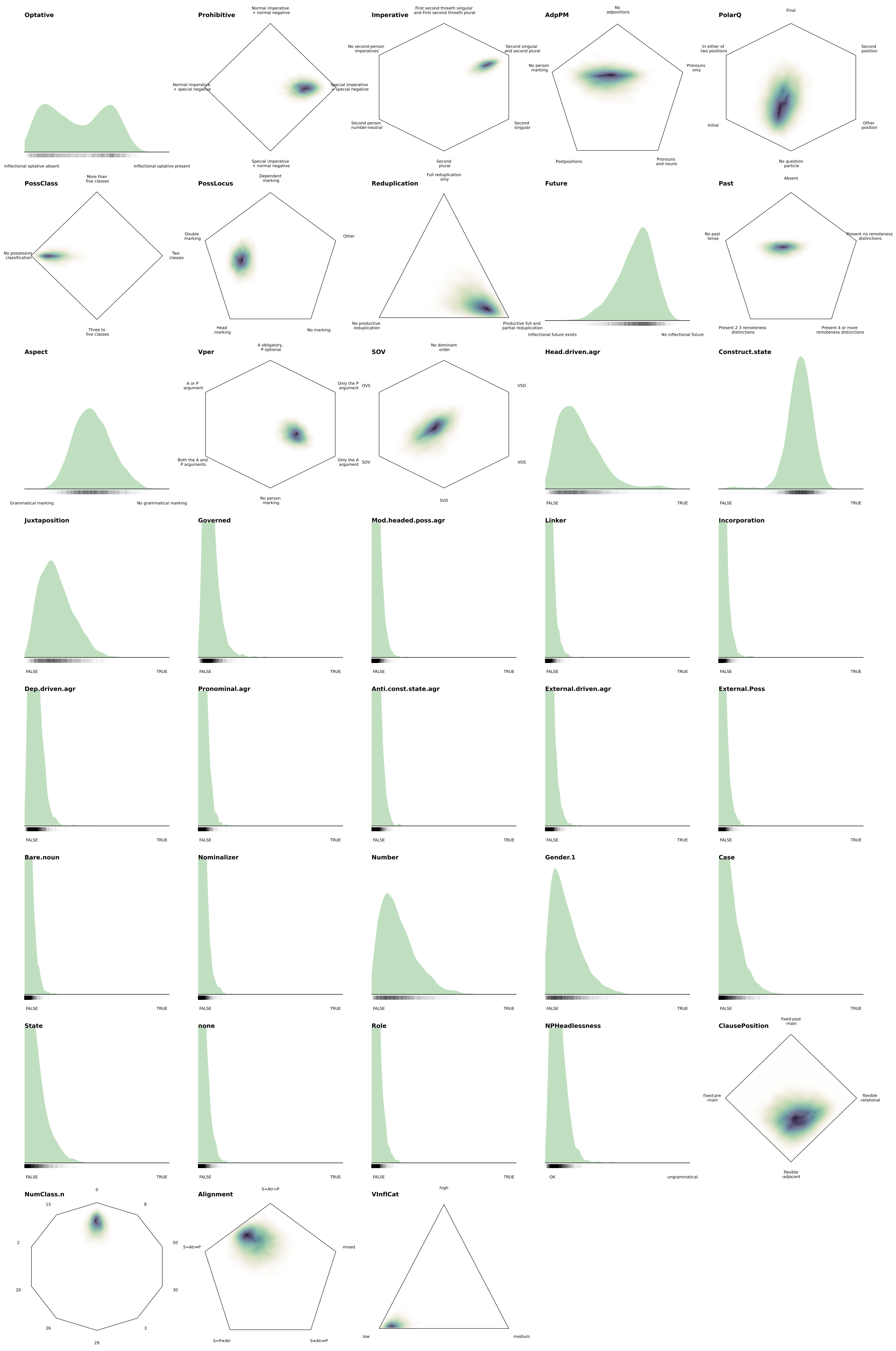

Figure S8: The state probabilities $\alpha$ for each feature in the universal distribution. (2/2)

###### S3.2.3 Areal distribution

The following pages show the state preferences $\gamma_{f,Z_1}$ for each features $f$ in the areal distribution of area $Z_1$.

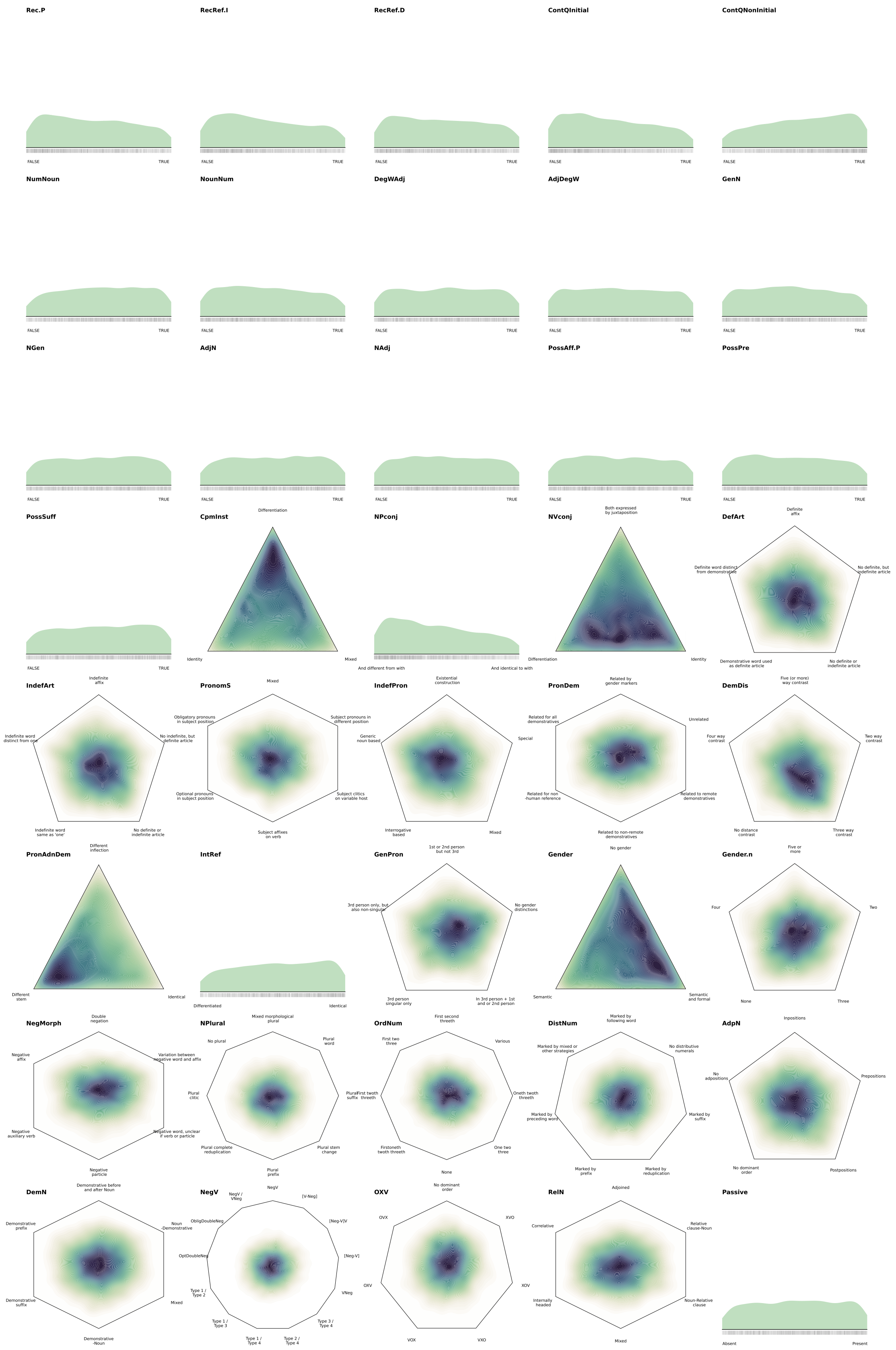


Figure S9: The state probabilities in the areal distribution $\gamma_{f,Z_1}$ for each feature $f$. (1/2)

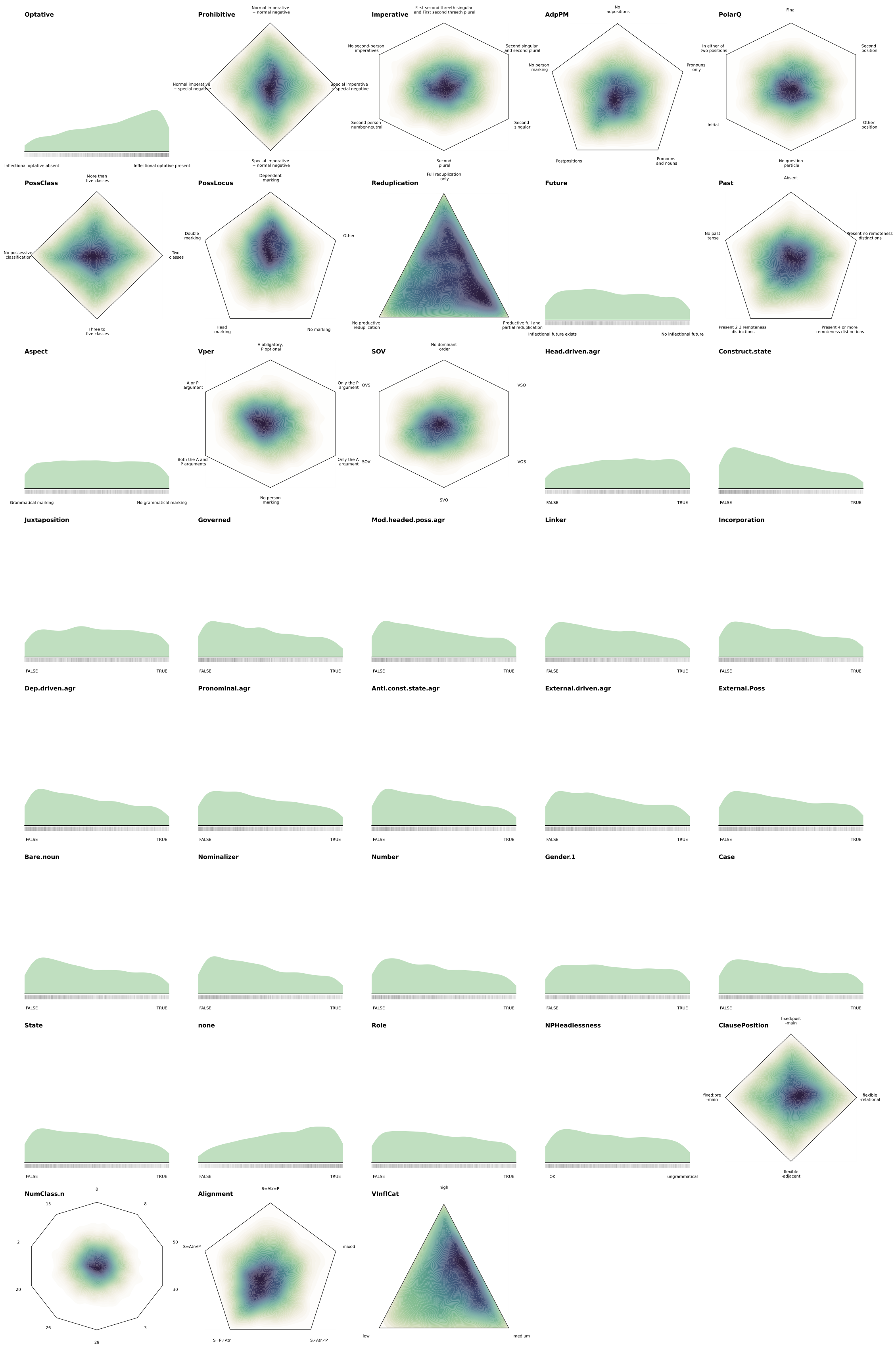

Figure S10: The state probabilities in the areal distribution $\gamma_{\cdot,Z_1}$ for each feature $f$ (2/2)

###### S3.2.4 Weights

The sBayes model defines the likelihood of each feature as a weighted sum over the three mixture components – universal, inheritance and cluster distribution. The mixture weights, $w$, indicate the influence of each mixture component on each feature $f$. The following pages show plots of the inferred weights per feature. The pink dots show the mean of the posterior distribution and the kernel density estimates in the background show the variation of all samples in the posterior distribution. The features are sorted by the strength of the cluster weight in the mean of the posterior distribution, that is, features that are more strongly affected by the cluster distribution are listed first.

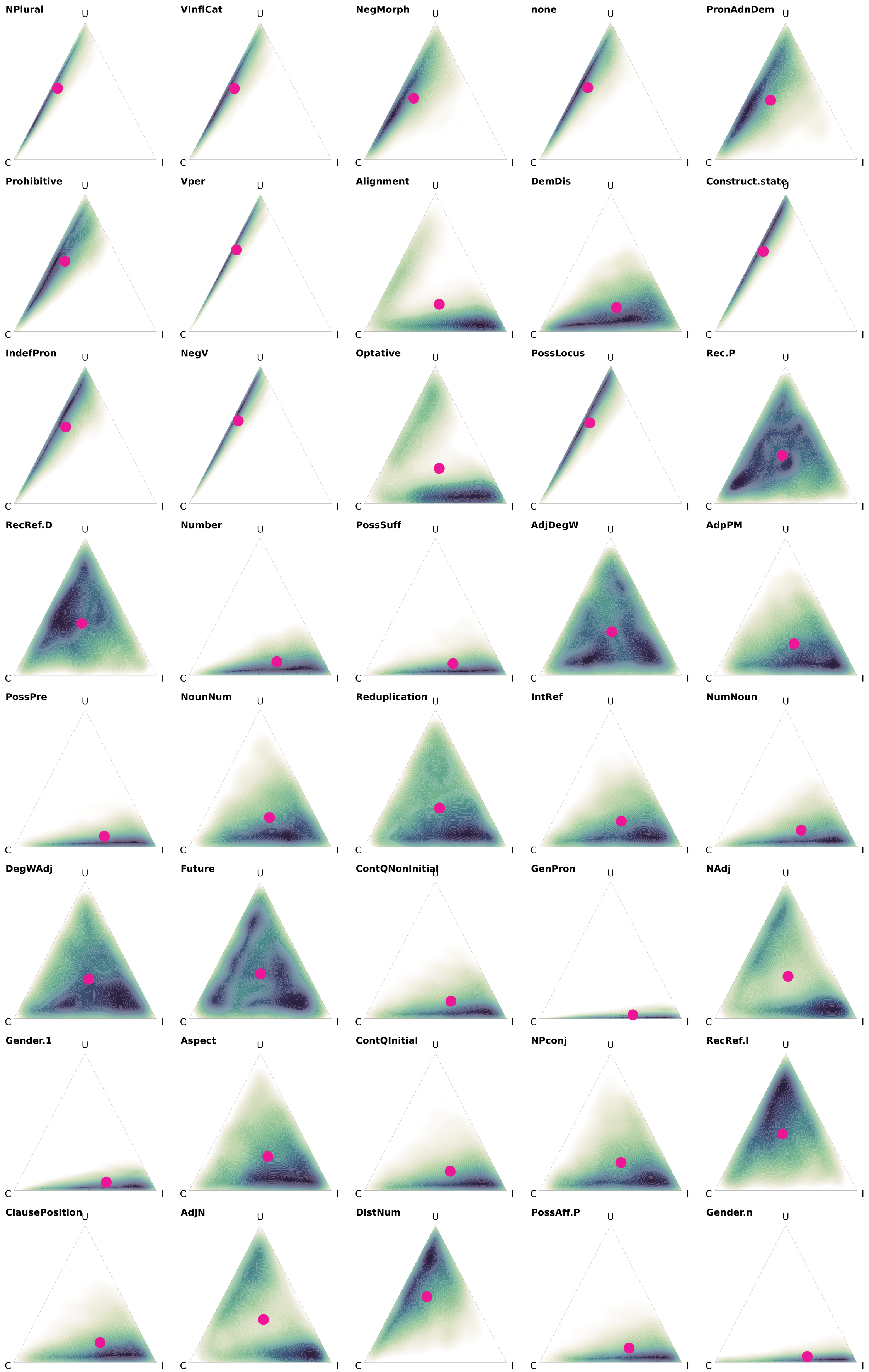


Figure S11: The mixture weights $w_{\cdot,f}$ for each feature $f$. (1/2)

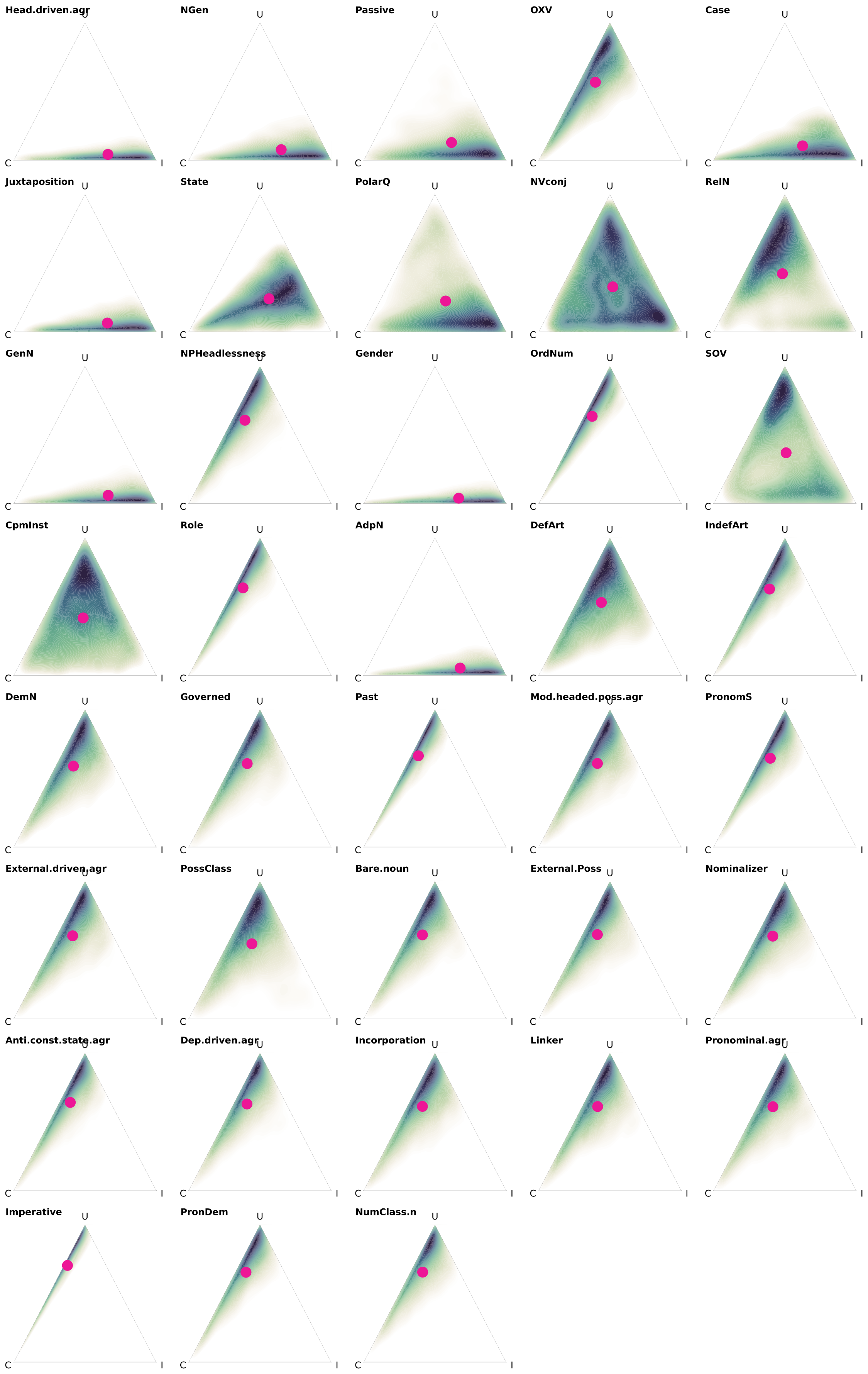


Figure S12: The mixture weights $w_{\cdot,f}$ for each feature $f$. (2/2)
